# Alzheimer’s-associated upregulation of mitochondria-associated ER membranes after traumatic brain injury

**DOI:** 10.1101/2020.11.13.381756

**Authors:** Rishi R. Agrawal, Delfina Larrea, Yimeng Xu, Lingyan Shi, Hylde Zirpoli, Leslie G. Cummins, Valentina Emmanuele, Donghui Song, Taekyung D. Yun, Frank P. Macaluso, Wei Min, Steven G. Kernie, Richard J. Deckelbaum, Estela Area-Gomez

**Author notes:** These authors contributed equally to this work. Please address correspondence to Rishi R. Agrawal and Estela Area-Gomez. Denali Therapeutics, 161 Oyster Point Blvd., South San Francisco, CA, 94080, USA. Centro de Investigaciones Biológicas Margarita Salas – CSIC, C. Ramiro de Maeztu, 9, 28040 Madrid, Spain (34-918-37-31-12).

## Abstract

Traumatic brain injury (TBI) can lead to neurodegenerative diseases such as Alzheimer’s disease (AD) through mechanisms that remain incompletely characterized. Similar to AD, TBI models present with cellular metabolic alterations and modulated cleavage of amyloid precursor protein (APP). Specifically, AD and TBI tissues display increases in amyloid-β as well as its precursor, APP C-terminal fragment of 99 a.a. (C99). Our recent data in cell models of AD indicate that C99, due to its affinity for cholesterol, induces the formation of transient lipid raft domains in the ER known as mitochondria-associated ER membranes (“MAM” domains). The formation of these domains recruits and activates specific lipid metabolic enzymes that regulate cellular cholesterol trafficking and sphingolipid turnover. Increased C99 levels in AD cell models promote MAM formation and significantly modulate cellular lipid homeostasis. Here, these phenotypes were recapitulated in the controlled cortical impact (CCI) model of TBI in adult mice. Specifically, the injured cortex and hippocampus displayed significant increases in C99 and MAM activity, as measured by phospholipid synthesis, sphingomyelinase activity and cholesterol turnover. In addition, our cell type-specific lipidomics analyses revealed significant changes in microglial lipid composition that are consistent with the observed alterations in MAM-resident enzymes. Altogether, we propose that alterations in the regulation of MAM and relevant lipid metabolic pathways could contribute to the epidemiological connection between TBI and AD.

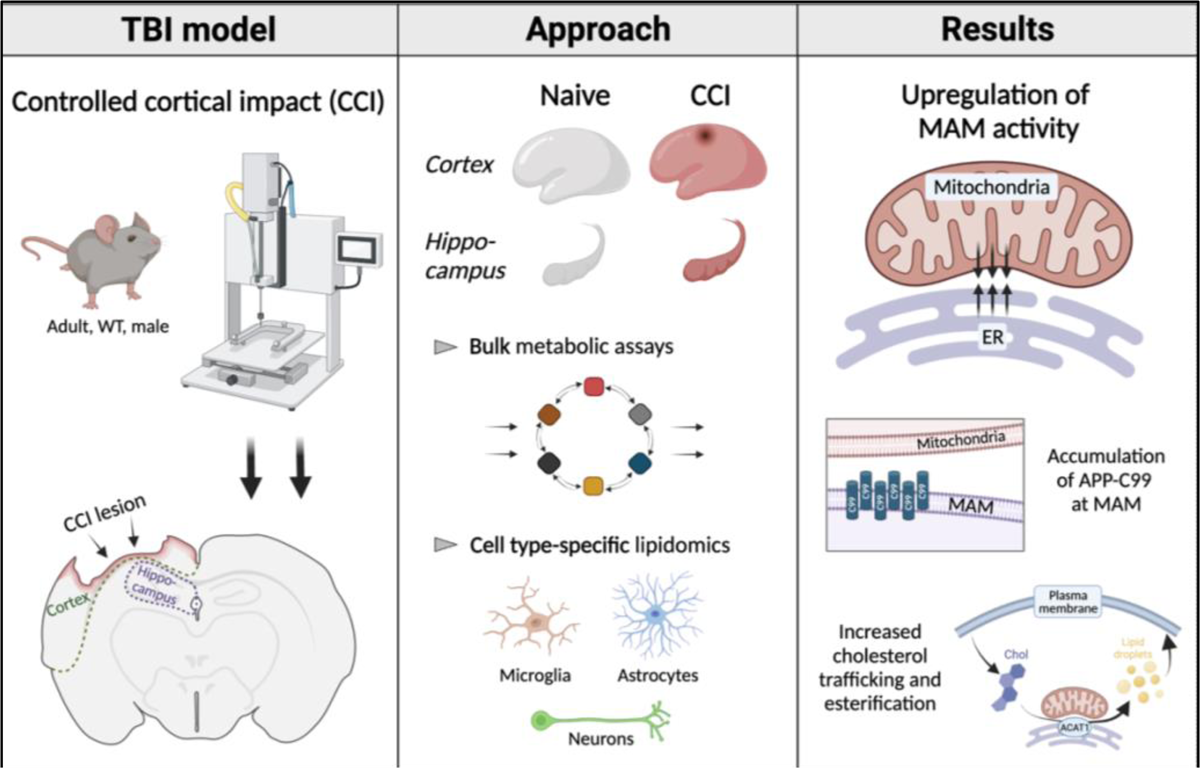

## Introduction

In the United States, traumatic brain injury (TBI) is the leading cause of death and disability for people under the age of 45 and incurs an annual economic cost of $76.5 billion. According to the Centers for Disease Control and Prevention (CDC), in 2014, 2.5 million people visited the emergency room due to a TBI, with 56,800 deaths; approximately 80,000 of these patients developed long-term disabilities (statistics sourced from the Brain Trauma Foundation and the CDC). Patients often fully recover after single TBI episodes, but a history of multiple TBIs is correlated with increased vulnerability to neurodegenerative diseases such as Alzheimer disease (AD) (Lye and Shores 2000; Van Den Heuvel et al 2007; McGuire 2018). Despite this epidemiological connection, the mechanism by which TBI can lead to AD-like neurodegeneration is unknown.

Our current understanding of the molecular pathogenesis of TBI largely derives from rodent studies that aim to recapitulate established AD signatures, and uncover common signaling aberrations. Indeed, it has been reported that AD models and brain injury models converge, at early stages, on brain lipid metabolic alterations, cell membrane damage, and organelle dysfunction (Adibhatla et al 2006; Chan et al 2012; Hiebert et al 2015; Montesinos et al 2020b). Moreover, cardinal AD features such as amyloid-β (Aβ) deposition, tau hyperphosphorylation, and neurofunctional deficits have been reported in TBI models (Washington et al 2012; Tsitsopoulos and Marklund 2013; Xu et al 2021). These studies suggest that TBI induces alterations in the cleavage of amyloid precursor protein (APP) similar to those found in AD (Ikonomovic et al 2004). Indeed, Aβ and tau deposition have been reported in human TBI tissues (Smith et al 2003; Uryu et al 2007; Gorgoraptis et al 2019), albeit inconsistently and sometimes manifesting in a minority of analyzed samples (Roberts et al 1991; Roberts et al 1994; Ikonomovic et al 2004). This has led the field to consider additional potential contributors to TBI and AD pathogenic mechanisms. Of note, elevations in Aβ precursor, the APP C-terminal fragment of 99 a.a. (C99), have also been reported in TBI (Chen et al 2004; Cartagena et al 2016) as well as in AD (Jiang et al 2010; Lauritzen et al 2012; Cavanagh et al 2013; Lauritzen et al 2016; Mondragón-Rodríguez et al 2018; Bourgeois et al 2018; Lauritzen et al 2019). While Aβ toxicity has been widely characterized, potential pathogenic effects of other APP fragments (such as C99) are not fully understood.

Recent data from our group indicates that C99, via its capacity to directly bind cholesterol (Beel et al 2010), can induce the formation of intracellular lipid rafts in the ER called mitochondria-associated ER membranes, or “MAM” domains (Montesinos et al 2020c). The formation of MAM domains modulates the lipid milieu of the ER and creates localized “signaling” platforms where specific enzymes are recruited for the concomitant regulation of numerous cellular pathways, including cellular lipid metabolism and membrane lipid composition (Vance 2014). Our data further show that, in AD models, elevations in C99 cause increased formation of MAM domains and the lipid metabolic functions regulated in these regions (Pera et al 2017). Specifically, at early disease stages, both familial (FAD) and sporadic (SAD) AD cells showed MAM-dependent increases in the turnover of sphingolipids and cholesterol, as well as significant alterations in the lipid composition of cell membranes (Area-Gomez et al 2012; Pera et al 2017; Montesinos et al 2020c). These early-stage findings in AD models prompted us to investigate whether similar pathologies are observable during the acute phase following TBI, as a potential early molecular link between TBI and AD.

In this preliminary study, using the controlled cortical impact (CCI) model of TBI in adult mice, we report upregulated function of MAM domains following a single, moderate injury. These defects in MAM regulation are associated with increased levels of C99 in MAM domains and the activation of cholesterol and sphingolipid turnover. These alterations modulate the lipidome of brain tissues and purified microglial, astrocytic and neuronal populations while preserving mitochondrial respiratory functionality. Through this groundwork analysis, we propose that TBI episodes could induce C99-mediated upregulation of MAM and the subsequent abrogation of lipid homeostasis, leading to AD-like molecular and cellular phenotypes.

## Materials and Methods

### Mouse husbandry

All animal husbandry was conducted in accordance with the Guide for the Care and Use of Laboratory Animals published by the National Institutes of Health. Specific procedures were approved by the Institutional Animal Care and Use Committee at Columbia University (protocols AC-AAAO5307 and AC-AAAY6450). Wild-type (WT) male, C57BL/6J mice (12-16 weeks in age) were ordered from Jackson labs and housed under a 12h light/12h dark cycle.

### Controlled cortical impact (CCI) injury

Twenty minutes following IP injection of general analgesic (5 mg/kg carprofen), the mouse was placed in an induction chamber and exposed to 4% isoflurane supplemented with oxygen at a flow rate of 1 liter per minute (LPM; gas supplied via an Isoflurane Vaporizer, Summit Anesthesia Solutions) for general anesthesia. Once breathing was stabilized and there was no response to toe pinch, head fur was clipped and the mouse was secured in a stereotaxic frame to initiate surgery. Anesthesia was maintained during surgery through a nose cone supplying isoflurane (eventually reduced to 2%) and oxygen (maintained at 1 LPM), verified every 2 min via lack of response to toe pinch. Body temperature was maintained at 37°C by keeping the mouse on a heating pad (Adroit Medical Systems) throughout the procedure. The scalp was disinfected by swabbing 3 times with fresh Prevantics antiseptic wipes (3.15% chlorhexidine gluconate/70% isopropanol, Henry Schein). Local analgesic (2 mg/kg bupivacaine) was injected subcutaneously at the injury site and 1 drop of Puralube ophthalmic ointment (Dechra Pharmaceutics) was applied to each eye. A midline incision was made to the scalp using a scalpel and a craniectomy was performed to create a 5 mm X 5 mm hole in the skull between the bregma and lambda and to the left of the sagittal suture. A single cortical contusion was made to the exposed dura mater using the Impact One impactor (Leica Biosystems) with the following injury parameters: 4.5 m/s velocity, 1.2 mm depth, 0.3 sec dwell time. Following injury, the incision was closed with sutures and Neosporin topical antibiotic ointment was applied. The mice were monitored during recovery from anesthesia. Following recovery, the mice were placed in post-procedure housing and monitored daily for wellness. General analgesic (5 mg/kg carprofen) was delivered via intraperitoneal (IP) injection 20 min before injury as well as 1 day (1d) and 2d after injury. Mice were sacrificed 1, 3 or 7 days after injury via cervical dislocation or transcardial perfusion.

### Cytochrome C oxidase (COX) activity staining

Mice were euthanized 1, 3 or 7 days after CCI (alongside age-matched naïve controls) by cervical dislocation. The brain was extracted and immediately snap-frozen in isopentane at −40°C for 35 sec. Following storage at −80°C, the brains were embedded in OCT medium (Thermo Fisher) and sectioned at 8 μm thickness using a CM3050S cryostat (Leica Biosystems). The sections were stored at −80°C until staining, at which point they were placed at room temperature (RT) for 25 min before staining to ensure adhesion to the slide. The staining solution consisted of 5.2 mM 3,3-diaminobenzidine (Millipore Sigma D8001), 0.09 mM cytochrome C (Millipore Sigma C2506), and 1.4 μM catalase (Millipore Sigma C1345) in 5 mM phosphate buffer, pH 7.4, filtered through Whatman paper. The stain was applied to the tissue for 25 min at 37°C in the dark, followed by 3 gentle and quick washes with ddH_2_O. The slides were coverslipped using warmed glycerin jelly as mounting medium. Images were collected the next day using a Nikon Eclipse 80i brightfield microscope with a 4X objective lens.

### Fluoro-Jade C (FJC) staining

Mice were euthanized 1, 3 or 7 days after CCI (alongside age-matched naïve controls) via transcardial perfusion with 40 mL 1X phosphate-buffered saline (PBS) followed by 80 mL 4% paraformaldehyde (PFA, Sigma) in PBS at a rate of 10 mL/min. The brain was removed and post-fixed in 4% PFA ON at 4°C with gentle agitation, cryo-protected in 30% sucrose for 2-3d at 4°C with gentle agitation (until the brains sank), snap-frozen in isopentane at −40°C for 35 sec, embedded in OCT medium (Thermo Fisher) and sectioned at a thickness of 16 μm using a CM3050S cryostat (Leica Biosystems, Buffalo Grove, IL). Following storage at −80°C, the slides were equilibrated at RT for 25 min and stained using the Fluoro-Jade C (FJC) Ready-to-Dilute Staining Kit (Biosensis TR-100) following manufacturer instructions. The slides were coverslipped using VECTASHIELD Antifade Mounting Medium (Vector Laboratories H-1000-10, Newark, CA). Imaging was conducted at room temperature using a confocal fluorescence microscope (Zeiss LSM710) with 10X and 20X objective lenses. Images were collected as Z-stacks (6 focal planes, 1 μm step) and exported in maximum projection. The images were converted to 8-bit images using ImageJ/Fiji, and FJC+ and DAPI+ cells were counted following the application of a thresholding algorithm. Quantification results are represented as percentage of DAPI+ cells that are also positive for FJC.

### Lipidomics analysis (bulk)

At the indicated timepoints after CCI, mice were euthanized (alongside age-matched naïve controls) via cervical dislocation and the brain was immediately extracted. The ipsilateral cortex and hippocampus (and corresponding regions in naïve animals) were microdissected on ice and homogenized in a Dounce homogenizer in Vance buffer (225 mM D-mannitol, 25 mM HEPES-KOH, 1 mM EGTA, pH 7.4) containing protease inhibitor (Millipore Sigma 11836170001). Protein concentration was measured using the Quick Start Bradford Protein Assay Kit 1 (Bio-Rad 5000201) and a Tecan Infinite F200 PRO spectrophotometer. Equal protein amounts of each sample (100 μg) were used for lipid extraction. Lipid extracts were prepared via chloroform–methanol extraction, spiked with appropriate internal standards, and analyzed using a 6490 Triple Quadrupole LC/MS system (Agilent Technologies) as previously described (Chan et al 2012). Free cholesterol and cholesteryl esters were separated with normal-phase HPLC using an Agilent Zorbax Rx-Sil column (inner diameter 2.1 Å∼ 100 mm) under the following conditions: mobile phase A (chloroform:methanol:1 M ammonium hydroxide, 89.9:10:0.1, v/v/v) and mobile phase B (chloroform:methanol:water:ammonium hydroxide, 55:39.9:5:0.1, v/v/v/v); 95% A for 2 min, linear gradient to 30% A over 18 min and held for 3 min, and linear gradient to 95% A over 2 min and held for 6 min. Quantification of lipid species was accomplished using multiple reaction monitoring (MRM) transitions that were developed in earlier studies (Chan et al 2012) in conjunction with referencing of appropriate internal standards. Values are represented as mole fraction with respect to total lipid (mole percentage). Lipid mass (in moles) of any specific lipid was normalized by the total mass (in moles) of all the lipids measured (Chan et al 2012).

### Magnetic-activated cell sorting (MACS) isolation of microglia, astrocytes and neurons

At the indicated timepoints after CCI, mice were euthanized by transcardial perfusion with 50 mL 1X Hanks’ Balanced Salt Solution ++ (HBSS with Ca^2+^ and Mg^2+^, without phenol red or Na_2_CO_3,_ Thermo Fisher 24020117), pH 7.4, at a rate of 10 mL/min. The brain was extracted and the ipsilateral cortex and hippocampus (and corresponding regions in naïve animals) were microdissected on ice. This step was not done in the validation experiments shown in Fig. 4**-10** as those were conducted in cells isolated from the whole brain (to generate enough starting material for the qPCR assays). For the remainder of the protocol, MACS+ Buffer (1X HBSS++, 15 mM HEPES and 0.5% BSA in ddH_2_O, pH 7.4) was the primary buffer in all steps. The tissue was dissociated using the Neural Tissue Dissociation Kit – Postnatal Neurons (Miltenyi Biotec 130-094-802) following manufacturer instructions with modifications: a 37°C oven with tube rotator was used instead of the GentleMACS dissociator, enzymes and buffer volumes were doubled (per gram of tissue), trituration was conducted with a trimmed P1000 low-retention pipet tip (Denville), and a 70-μm Falcon cell strainer (Fisher Scientific 08-771-2) was used instead of the MACS SmartStrainer. For the validation qPCR experiments, a 200 μL aliquot of this suspension was taken as the “unsorted” sample.

For myelin removal, Myelin Removal Beads II (Miltenyi Biotec 130-096-731) were used. Per 0.5g starting material weight, the pellet from dissociation was resuspended in 3.6 mL MACS+ buffer to which 400 μL beads was added. The sample was incubated at 4°C for 15 min with agitation every 5 min. The reaction was suspended by adding 5 volumes MACS+ buffer, mixing by inversion and pelleting of the cells at 300x*g* for 10 min at 4°C. The pellet was resuspended in 6 mL MACS+ buffer and evenly distributed between two LS columns (Miltenyi Biotec 130-042-401) in the presence of a magnetic field (generated by the QuadroMACS magnetic separator, Miltenyi Biotec). After washing the columns with 3 mL MACS+ buffer three times, the flow-through was centrifuged at 300x*g* for 10 min at 4°C.

For microglia isolation, the resultant pellet was resuspended in 875 μL MACS+ buffer in a 5-mL epitube, to which 125 μL CD11b microbeads (Miltenyi Biotec 130-049-601) was added. The sample was placed on a rotating wheel at 4°C for 30 min, after which the tube was filled with buffer and centrifuged at 300x*g* for 10 min at 4°C. The pellet was resuspended in 4 mL MACS+ buffer and divided evenly between two LS columns in the presence of a magnetic field. The columns were washed 3 times each with 3 mL MACS+ buffer, and the total flow-through was set aside for astrocyte isolation. The microglia were collected by removing the columns from the magnetic field, applying 5 mL MACS+ buffer, and using the plunger to elute the captured cells. Both the eluent and flow-through were centrifuged at 300x*g* for 10 min at 4°C.

For astrocyte isolation, the pellet of the flow-through was resuspended in 700 μL MACS+ buffer, to which 150 μL FcR Blocking Reagent (Miltenyi Biotec 130-092-575) was added. The sample was placed on a rotating wheel at 4°C for 30 min, followed by addition of 150 μL Anti-ACSA-2 microbeads (Miltenyi Biotec 130-097-678). The sample was placed on a rotating wheel at 4°C for 30 min, after which the tube was filled with buffer and centrifuged at 300x*g* for 10 min at 4°C. The pellet was resuspended in 4 mL MACS+ buffer and divided evenly between two LS columns in the presence of a magnetic field. The columns were washed 3 times each with 3 mL MACS+ buffer, and the total flow-through was set aside as the “neurons.” The astrocytes were collected by removing the columns from the magnetic field, applying 5 mL MACS+ buffer, and using the plunger to elute the captured cells. Both the eluent and flow-through were centrifuged at 300x*g* for 10 min at 4°C. The supernatants were removed and the pellets were stored for downstream applications.

### Lipidomics analysis (MACS-sorted populations)

Following MACS isolation of cell type-specific populations, the final pellets were snap-frozen on dry ice and stored at −80°C until lipid extraction. Lipidomics analysis was conducted as described above in the “Lipidomics Analysis (bulk)” section; however, since protein concentration was not determined (due to the high concentration of BSA in the isolation buffer), quantities (moles) of the extracted lipid species were normalized by quantity (moles) of total lipid extracted to determine mole percentage.

### Mitochondrial DNA (mtDNA) to nuclear DNA (nDNA) ratio

Mice were euthanized 1, 3 or 7 days after CCI (alongside age-matched naïve controls) by cervical dislocation. The brain was extracted and the ipsilateral cortex and hippocampus (and corresponding regions in naïve animals) were microdissected on ice. The tissues were homogenized in a Dounce homogenizer in Vance buffer (225 mM D-mannitol, 25 mM HEPES-KOH, 1 mM EGTA, pH 7.4) containing protease inhibitor (Millipore Sigma 11836170001). Protein concentration was measured using the Quick Start Bradford Protein Assay Kit 1 (Bio-Rad 5000201) in a Tecan Infinite F200 PRO spectrophotometer, and equal protein amounts (200 μg) were used for each sample. Following digestion with 1 mg/mL proteinase K (Millipore Sigma P4850), DNA was extracted using a standard phenol-chloroform extraction protocol with isopropanol for precipitation and ethanol for cleaning. RNA was digested using RNase A/T1 mix (Thermo Fisher EN0551) and DNA concentration was measured via Nanodrop 1000. Quantitative PCR was conducted under standard conditions using the reaction composition in Table 1.

**Table 1:**
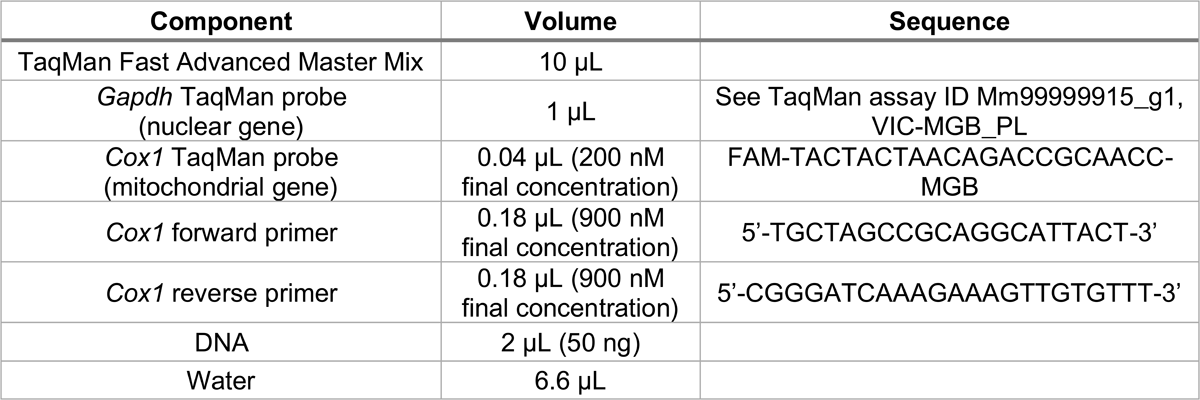
Composition of mtDNA:nDNA ratio qPCR reaction.

### Oil Red O staining

Mice were euthanized 3d after CCI (alongside age-matched naïve controls) via transcardial perfusion with 40 mL PBS followed by 80 mL 4% PFA (in PBS) at 10 mL/min. The brain was removed and post-fixed in 4% PFA ON at 4°C with gentle agitation, cryo-protected in 30% sucrose for 2-3d at 4°C with gentle agitation (until the brains sank), snap-frozen in isopentane at −40°C for 35 sec, embedded in OCT medium (Thermo Fisher) and sectioned at a thickness of 16 μm using a CM3050S cryostat (Leica Biosystems, Buffalo Grove, IL). Following storage at −80°C, the slides were equilibrated at RT for 25 min and placed in a humidified chamber. The slides were incubated with 10% formalin for 1 min, dipped in ddH_2_O for 30 sec and allowed to air-dry. In the humidified chamber, Oil Red O 0.5% solution in propylene glycol (Poly Scientific R&D Corp. s1848) was applied for 1h, followed by three 5-min dips in ddH_2_O. The slides were air-dried and coverslipped with Fluoromount-G (Thermo Fisher 00-4958-02). Images were immediately collected with a Nikon Eclipse 80i brightfield microscope using a 40X objective.

### Quantitative polymerase chain reaction (qPCR; bulk)

Mice were euthanized 3d after CCI (alongside age-matched naïve controls) and the brain was immediately extracted. The ipsilateral cortex and hippocampus (and corresponding regions in naïve animals) were microdissected on ice and immediately homogenized by pipetting and vortexing in TRIzol (Thermo Fisher). Total RNA was extracted using manufacturer instructions and tested for purity (by measuring A_260_/A_280_ and A_260_/A_230_ ratios via a Nanodrop 1000 Thermo Fisher) and for integrity (by agarose gel electrophoresis and detection of bands corresponding to 28S and 18S ribosomal RNA). Following digestion with RQ1 RNase-free DNase (Promega M6101), reverse transcription was performed with the High Capacity cDNA Reverse Transcription Kit (Thermo Fisher 4368814). Quantitative PCR was conducted in triplicate in a StepOne Plus real-time PCR machine (Applied Biosystems) using TaqMan Fast Advanced Master Mix (Thermo Fisher 4444556). The expression of each gene was analyzed using pre-designed TaqMan Probes (Thermo Fisher), with *Gapdh* (assay ID Mm99999915_g1, conjugated to the VIC-MGB_PL dye) serving as the housekeeping gene. The assay IDs of the probes used for the experimental genes tested, all conjugated to the FAM-MGB dye, are listed in Table 2. Data was analyzed using the ΔΔCt method.

**Table 2:** TaqMan probes used for bulk qPCR experiments.

| **Gene** | **Assay ID** |
| --- | --- |
| *Abca1* | Mm00442646_m1 |
| *Abcg1* | Mm00437390_m1 |
| *Acaca* | Mm01304257_m1 |
| *Aif1* | Mm00479862_g1 |
| *Bace1* | Mm00478664_m1 |
| *Cd36* | Mm00432403_m1 |
| *Cldn11* | Mm00500915_m1 |
| *Cpt1a* | Mm01231183_m1 |
| *Fasn* | Mm00662319_m1 |
| *Gfap* | Mm01253033_m1 |
| *Hif1a* | Mm00468869_m1 |
| *Ldlr* | Mm01177349_m1 |
| *Lrp1* | Mm00464608_m1 |
| *Npc1* | Mm00435300_m1 |
| *Pdk1* | Mm00554300_m1 |
| *Ppargc1a* | Mm01208835_m1 |
| *Scarb1* | Mm00450234_m1 |
| *Smpd1* | Mm00488319_g1 |
| *Smpd2* | Mm01188195_g1 |
| *Smpd3* | Mm00491359_m1 |
| *Smpd4* | Mm00547173_m1 |
| *Smpd5* | Mm01205829_g1 |
| *Srebf1* | Mm00550338_m1 |

### qPCR for cell type-specific markers in MACS-sorted populations

Following collection of pellets, total RNA was immediately extracted using the RNeasy Plus Micro Kit (Qiagen). Lysis was conducted by vortexing the pellet in 350 μL Buffer RLT Plus containing β-mercaptoethanol (BME) for 1 min. No carrier RNA was used. Purity was determined by measuring A_260_/A_280_ and A_260_/A_230_ ratios via a Nanodrop 1000. Because the RNeasy Plus kit includes gDNA eliminator columns, DNase digestion was not conducted as a separate step. Reverse transcription was performed using the High Capacity cDNA Reverse Transcription Kit (Thermo Fisher 4368814). Quantitative PCR was conducted in triplicate with TaqMan Fast Advanced Master Mix (Thermo Fisher 4444556) in a StepOne Plus real-time PCR machine (Thermo Fisher). The expression of each gene under study was analyzed using pre-designed TaqMan Probes, with *Rn18s/Rn45s* (encoding 18S rRNA and 45S pre-rRNA; assay ID Mm03928990_g1, conjugated to VIC-MGB_PL) serving as the housekeeping gene. The assay IDs of the experimental genes tested are listed in Table 3, with all probes conjugated to the FAM-MGB dye. Data from each mouse was analyzed separately. Grubbs’ test (⍺=0.05) was used to identify outliers, which were then removed.

**Table 3:** TaqMan probes used for qPCR experiments in MACS-sorted populations.

| Cell type | Marker gene | Assay ID |
| --- | --- | --- |
| Astrocytes | *Gfap* | Mm01253033_m1 |
|  | *S100b* | Mm00485897_m1 |
|  | *Slc1a2* | Mm01275814_m1 |
| Microglia | *Aif1* | Mm00479862_g1 |
|  | *Cx3cr1* | Mm02620111_s1 |
|  | *Itgam* | Mm00434455_m1 |
| Neurons | *Rbfox3* | Mm01248771_m1 |
|  | *Tubb3* | Mm00727586_s1 |
|  | *Dcx* | Mm00438400_m1 |
| Oligodendrocytes | *Cldn11* | Mm00500915_m1 |

### Phospholipid transfer assay

This assay was conducted as described previously (Area-Gomez 2014; Montesinos et al 2020a). Briefly, mice were euthanized 3d after CCI (alongside age-matched naïve controls) by cervical dislocation. The brain was extracted and the ipsilateral cortex and hippocampus (and corresponding regions in naïve animals) were microdissected on ice. The tissues were homogenized with a Dounce homogenizer in Vance buffer (225 mM D-mannitol, 25 mM HEPES-KOH, 1 mM EGTA, pH 7.4) containing protease inhibitor (Millipore Sigma 11836170001). Crude mitochondria fractions were prepared as described previously (Area-Gomez 2014; Montesinos and Area-Gomez 2020). Protein concentration was measured using the Quick Start Bradford Protein Assay Kit 1 (Bio-Rad 5000201) in a Tecan Infinite F200 PRO spectrophotometer, and 100 μg protein was brought to 50 μL in phospholipid transfer assay buffer (10 mM CaCl_2_, 25 mM HEPES-KOH, 0.01% Triton X-100) spiked with 0.04 mM ^3^H-serine (Perkin Elmer NET248005MC). The reactions were incubated at 37°C for 45 min and stored at −20°C as needed. Lipids were extracted and resolved through thin layer chromatography (TLC), followed by excision of relevant spots and scintillation counting as described previously (Area-Gomez 2014; Montesinos et al 2020a). For each experiment, the average of each species in the CCI group was divided by the average in the naïve group to calculate fold change of CCI over naïve. These numbers are represented in the graph and were used for statistical analysis.

### Seahorse analysis of mitochondrial respiration

Mitochondrial oxygen consumption rate (OCR) was determined as previously described (Agrawal et al 2020). In summary, mice were euthanized 1, 3 or 7 days after CCI (alongside age-matched naïve controls) by cervical dislocation. The brain was extracted and the ipsilateral cortex and hippocampus (and corresponding regions in naïve animals) were microdissected on ice. The tissues were homogenized in ∼10 volumes homogenization buffer (210 mM mannitol, 70 mM sucrose, 5 mM HEPES and 1 mM EDTA, pH 7.2 adjusted with KOH) using an appropriately sized Dounce homogenizer with a teflon pestle. An equal volume of washing buffer (homogenization buffer with 0.5% fatty acid-free BSA added) was used to wash the homogenizer and dilute the homogenate 2X. The total homogenate was centrifuged at 900x*g* for 10 min at 4°C. The supernatant was centrifuged at 900x*g* for 10 min at 4°C again to pellet any residual debris, and the resulting supernatant was centrifuged at 8000x*g* for 10 min at 4°C to pellet the crude mitochondria (CM) fraction. The pellets were resuspended in homogenization buffer (lacking BSA) and centrifuged again at 8000x*g* for 10 min at 4°C. The final pellet was resuspended in a small volume of homogenization buffer, and the protein concentration was measured using the Quick Start Bradford Protein Assay Kit 1 (Bio-Rad 5000201) in a Tecan Infinite F200 PRO spectrophotometer. For C-I experiments, 8 μg protein in a volume of 50 μL was added to each well; for C-II analysis, 6 μg protein in a volume of 50 μL was added to each well. The buffer used to further dilute the CM suspensions was assay buffer (220 mM mannitol, 70 mM sucrose, 5 mM KH_2_PO_4_, 5 mM MgCl_2_, 2 mM HEPES, 1 mM EGTA, 0.2% fatty acid-free BSA, pH 7.4 adjusted with KOH) with 5 mM pyruvate and 5 mM malate added for C-I assays, and 5 mM succinate and 2 μM rotenone added for C-II assays. The plate was centrifuged at 2000x*g* for 5 min at 4°C, and 400 μL additional buffer was added to each well (with 450 μL buffer added to blank wells). Oxygen consumption was measured at States 2, 3, 4, and 3-uncoupled after sequential addition of 3 mM ADP, 3.4 μM oligomycin, 5.7 μM FCCP and 3.4 μM Antimycin A (final concentrations, ∼10X were stocks added to the cartridge lid injection ports in advance), respectively.

### Stimulated Raman Scattering (SRS) imaging

Mice were euthanized 7d after CCI (alongside age-matched naïve controls) via transcardial perfusion with 40 mL PBS followed by 80 mL 4% PFA (in PBS) at a rate of 10 mL/min. The brain was removed and post-fixed in 4% PFA ON at 4°C with gentle agitation. The brains were embedded in 3% agarose gel in a cubic cryomold and coronally sectioned at 50 μm thickness using a VT1000S vibrating blade microtome (Leica Biosystems, Buffalo Grove, IL). The sections were rinsed in PBS ON at 4°C, mounted onto slides and coverslipped.

Imaging was conducted using an inverted laser-scanning microscope (FV1200, Olympus) optimized for near-IR throughput and a 25X water objective (XLPlanN, 1.05 N.A., MP, Olympus) with high near-IR transmission for SRS imaging. A picoEMERALD system (Applied Physics & Electronics) supplied a synchronized pulse pump beam (with tunable 720-990 nm wavelength, 5-6 ps pulse width, and 80-MHz repetition rate) and a Stokes beam (with fixed wavelength at 1064 nm, 6 ps pulse width, and 80 MHz repetition rate). Stokes was modulated at 8 MHz by an electronic optic modulator. Transmission of the forward-going pump and Stokes beams after passing through the samples was collected by a high N.A. oil condenser (N.A. = 1.4). A high O.D. bandpass filter (890/220, Chroma) was used to block the Stokes beam completely and to transmit only the pump beam onto a large area Si photodiode for detection of the stimulated Raman loss signal. The output current from the photodiode was terminated, filtered, and demodulated by a lock-in amplifier (Zurich, HF2LI) at 8 MHz to ensure shot-noise-limited detection sensitivity. The demodulated signal was fed into the analog channel of the FV1200 software FluoView 4.1a (Olympus) to form the image during laser scanning at a rate of 100 μs per pixel. For multi-channel SRS imaging, the pump wavelength was tuned so that the energy difference between pump and Stokes matched with the vibrational frequency as previously described (Shi et al 2018). λ_pump_= 1/(1/1064+10^-7^*v) where v is the vibrational frequency in cm^-1^. CH channels were acquired at 2845 cm^-1^ and 2940 cm^-1^ and unmixed as previously described (Shi et al 2018).

### Sphingomyelinase (SMase) assay

Mice were euthanized 3d after CCI (alongside age-matched naïve controls) by cervical dislocation. The brain was extracted and the ipsilateral hippocampus (and the corresponding region in naïve animals) was microdissected on ice. The tissues were homogenized in a Dounce homogenizer in Vance buffer (225 mM D-mannitol, 25 mM HEPES-KOH, 1 mM EGTA, pH 7.4) containing protease inhibitor (Millipore Sigma 11836170001). SMase activity was assayed as previously described (Pera et al 2017). Following measurement of the protein concentration using the Quick Start Bradford Protein Assay Kit 1 (Bio-Rad 5000201) in a Tecan Infinite F200 PRO spectrophotometer, 100 µg protein was assayed in a buffer containing 100 mM tris/glycine (pH 7.4), 1.55 mM Triton X-100, 0.025% fatty acid-free BSA, 1 mM MgCl_2_, and 400 µM bovine brain sphingomyelin (SM) spiked with 22,000 dpm ^3^H-SM (American Radiolabeled Chemicals ART 0481-50 µCi) at a final concentration of 1 nCi/sample. Reactions were incubated at 37°C overnight, followed by quenching with 1.2 mL ice-cold 10% trichloroacetic acid, incubation at 4°C for 30 min, and centrifugation at 1500x*g* at 4°C for 20 min. 1 mL supernatant was transferred to clean tubes, 1 mL ether was added, and the mixture was vortexed and centrifuged at 1500x*g* for 5 min at 4°C. 800 µL of the bottom phase was transferred to scintillation vials containing 5 mL ScintiVerse BD Cocktail (Fisher Scientific SX18-4). The vials were vortexed and radioactivity was measured in a Scintillation Counter (Tri-Carb 2819TR, Perkin Elmer).

### Subcellular fractionation

Mice were euthanized 3d after CCI (alongside age-matched naïve controls) via cervical dislocation and the brain was immediately extracted. The ipsilateral cortex and hippocampus (and corresponding regions in naïve animals) were microdissected on ice and homogenized in a Dounce homogenizer in Vance buffer (225 mM D-mannitol, 25 mM HEPES-KOH, 1 mM EGTA, pH 7.4) containing protease inhibitor (Millipore Sigma 11836170001). Subcellular fractionation was conducted as described previously (Montesinos and Area-Gomez 2020).

### Succinate dehydrogenase (SDH) activity staining

Mice were euthanized 1, 3 or 7 days after CCI (alongside age-matched naïve controls) by cervical dislocation. The brain was extracted and immediately snap-frozen in isopentane at −40°C for 35 sec. Following storage at −80°C, the brains were sectioned at 8 μm thickness using a CM3050S cryostat (Leica Biosystems). The sections were stored at −80°C until staining, at which point they were placed at RT for 25 min before staining to ensure adhesion to the slide. The staining solution consisted of 5 mM EDTA, 1 mM KCN, 0.2 mM phenazine methosulfate, 50 mM succinic acid and 1.5 mM Nitro Blue in 5 mM phosphate buffer, pH 7.6, filtered through Whatman paper. The stain was applied for 10 min at 37°C in the dark, followed by 3 gentle and quick washes with ddH_2_O. The slides were coverslipped using warmed glycerin jelly as mounting medium. Images were collected the next day with a Nikon Eclipse 80i brightfield microscope with a 4X objective lens.

### Transmission electron microscopy

Mice were euthanized 3d after CCI (alongside age-matched naïve controls) via transcardial perfusion with 40 mL PBS followed by 80 mL 4% PFA (in PBS) at a rate of 10 mL/min. The brain was removed, post-fixed in 4% PFA ON at 4°C with gentle agitation, and sectioned at a thickness of 100 μm using a VT1000S vibrating blade microtome (Leica Biosystems). The sections were fixed by gentle agitation for 1h at RT with 2% PFA and 2.5% glutaraldehyde in 0.1 M sodium cacodylate buffer. The sections were then post-fixed with 1% osmium tetroxide followed by 2% uranyl acetate, dehydrated through a graded series of ethanol, and sandwiched between Aclar sheets for embedding in LX112 resin (LADD Research Industries). The area of interest was extracted, glued to a blank resin block and ultrathin (80 nm) sections were cut on a Leica EM Ultracut UC7, stained with uranyl acetate followed by lead citrate and viewed on a JEOL 1400 Plus transmission electron microscope at 80kv.

### Western blot for APP-C99

Following subcellular fractionation (described above), protein concentration was measured using the Quick Start Bradford Protein Assay Kit 1 (Bio-Rad 5000201) in a Tecan Infinite F200 PRO spectrophotometer. For each fraction, 10 μg protein was combined with 1 μL 10% Triton X-100 (Sigma) and brought to a volume of 33 μL in Vance buffer (225 mM D-mannitol, 25 mM HEPES-KOH, 1 mM EGTA, pH 7.4). The ratio of 10 μg Triton:1 μg protein was maintained. The sample was vortexed for 15 sec, incubated on ice for 1h with occasional short vortexes, and combined with 11 μL loading buffer (4X NuPAGE LDS sample buffer, Thermo Fisher, containing 10% β-mercaptoethanol). The sample was heated at 95°C for 5 min and loaded into a 4-12% Criterion XT Bis-Tris Protein Gel, 12+2 well, midi format (Bio-Rad 3450123) using 1X XT MES Running Buffer (Bio-Rad 1610789). Electrophoresis was conducted at 80-120V. The sample was transferred to an Immuno-Blot PVDF membrane (Bio-Rad 1620177) in standard tris-glycine transfer buffer with 20% methanol and 0.04% SDS at 150 mA for 90 min in a wet transfer. The primary antibodies used are as follows: APP-C99 [Biolegend 805701, clone M3.2 (Morales-Corraliza et al 2009)]; APP C-terminus [Sigma A8717, detects both C99 and C83 (Pera et al 2017)]; Erlin-2 [Abcam ab129207 (Liu et al 2020)] and TOM20 [Santa Cruz sc-11415 (Rodriguez-Sinovas et al 2006)]. The secondary antibodies (linked to horseradish peroxidase) are: anti-rabbit IgG (Fisher Scientific 45-000-682) and anti-mouse IgG (Fisher Scientific 45-000-679). Band densitometry was conducted using Fiji.

### Western blot for TOM20 and OxPhos complexes

Mice were euthanized 1, 3 or 7 days after CCI (alongside age-matched naïve controls) by cervical dislocation. The brain was extracted and the ipsilateral cortex and hippocampus (and corresponding regions in age-matched naïve animals) were microdissected on ice. The tissues were homogenized in a Dounce homogenizer in Vance buffer (225 mM D-mannitol, 25 mM HEPES-KOH, 1 mM EGTA, pH 7.4) containing protease inhibitor (Millipore Sigma 11836170001). Protein concentration was measured using the Quick Start Bradford Protein Assay Kit 1 (Bio-Rad 5000201) in a Tecan Infinite F200 PRO spectrophotometer. 10 μg protein was combined with 4X Laemmli loading buffer in a final volume of 20 μL, heated at 50°C, and loaded onto a Novex WedgeWell 4-20% tris-glycine mini 12-well SDS-PAGE gel. Electrophoresis was conducted at 80-120V using standard tris-glycine running buffer. The sample was transferred to an Immuno-Blot PVDF membrane (Bio-Rad 1620177) in standard tris-glycine transfer buffer with 20% methanol and 0.04% SDS at 150 mA for 2h in a wet transfer. The primary antibodies used are as follows: Total OxPhos Rodent WB antibody cocktail [Abcam ab110413 (Li et al 2009)]; TOM20 [Santa Cruz sc-11415 (Rodriguez-Sinovas et al 2006)]; β-actin [Sigma A5441 (Bunnell et al 2011)]; and Vinculin [Sigma V4505 (Sydor et al 1996)]. The secondary antibodies (linked to horseradish peroxidase) are: anti-rabbit IgG (Fisher Scientific 45-000-682) and anti-mouse IgG (Fisher Scientific 45-000-679). Band densitometry was conducted using Fiji.

### Statistical analysis

Data represent mean ± SD. All averages are the result of three or more separate animals, as indicated in the figure legends, processed independently. Historical and preliminary data were used to ensure adequate power and to limit the background associated with each assay (SigmaPlot 12.0). Data distribution was assumed to be normal. For all assays, statistical comparisons were made to naïve (uninjured) tissues from the same region and/or cell type harvested and assayed alongside CCI tissues. Statistical analyses were conducted using Microsoft Excel and GraphPad Prism v9.0.0. The authors were not blinded to the experimental groups while performing statistical analysis. Statistical significance was determined via two-tailed T-test or Ordinary one-way ANOVA followed by Bonferroni’s multiple comparisons test, as indicated in the figure legends. The significance level was set to ⍺=0.05 for all tests, and p-values below 0.05 were considered statistically significant.

## Results

### The controlled cortical impact (CCI) model of TBI induces region-specific cell death and glial activation

The experimental TBI models employed in pre-clinical research recapitulate features of human TBI to varying degrees (Ma et al 2019). In this study, we applied the controlled cortical impact (CCI) model to adult male wild-type (WT) C57BL/6J mice. During a CCI injury, a rapidly accelerating rod penetrates the cerebral cortex through an opening in the skull (Osier and Dixon 2016). We selected this model on the basis of its ability to recapitulate human TBI features in a reproducible manner (Bryan et al 1995; Washington et al 2012) and the ease with which injury parameters can be controlled (Osier and Dixon 2016). We performed an injury of moderate severity where the cortex is physically impacted but subcortical structures, including the hippocampus, are not (Osier and Dixon 2016).

We began this study by assaying known responses to brain injury to determine the regional specificity of activated molecular pathways. Our studies were conducted in the ipsilateral cortex (directly injured) and ipsilateral hippocampus (directly below the injury), and results were compared to age-matched naïve (uninjured) samples. We collected tissues 1, 3, and/or 7 days after injury because the APP and lipid metabolic phenotypes relevant to this study, whereby APP-C99 accumulation at MAM domains regulates cellular cholesterol and sphingolipid trafficking (Pera et al 2017; Montesinos et al 2020c), have been previously reported at these timepoints (Chen et al 2004; Roux et al 2016). Further, we sought to generate insight into whether these parameters were transient or sustained during the acute phase after injury.

A well-characterized phenotype of the CCI model is cell death, considered a major contributor to behavioral alterations after TBI (Girgis et al 2016). Since the extent and distribution of cell death differs based on injury parameters, we characterized this response in our model through FluoroJade-C staining, which labels degenerating neurons (Schmued et al 2005). We observed widespread cell death in the cortex at 1d and 3d that diminished by 7d after injury. Conversely, cell death was nearly absent in the hippocampus throughout the assayed post-injury time-course (Fig. 1A-C). Despite these differences in the levels of neuronal death, the activation of astrocytes and microglia, as measured by gene expression of *Gfap* [an astrocyte marker (Eng and Ghirnikar 1994)] and *Aif1* [a microglia marker (Postler et al 2000)], was observable in both brain regions (Fig. 1D/E). Oligodendrocyte activation, represented by gene expression of oligodendrocyte marker *Cldn11* (Bronstein et al 2000), was limited to the cortex (Fig. 1F). Altogether, our results demonstrate robust glial activation in the hippocampus in the absence of significant cell death.

**Figure 1:**
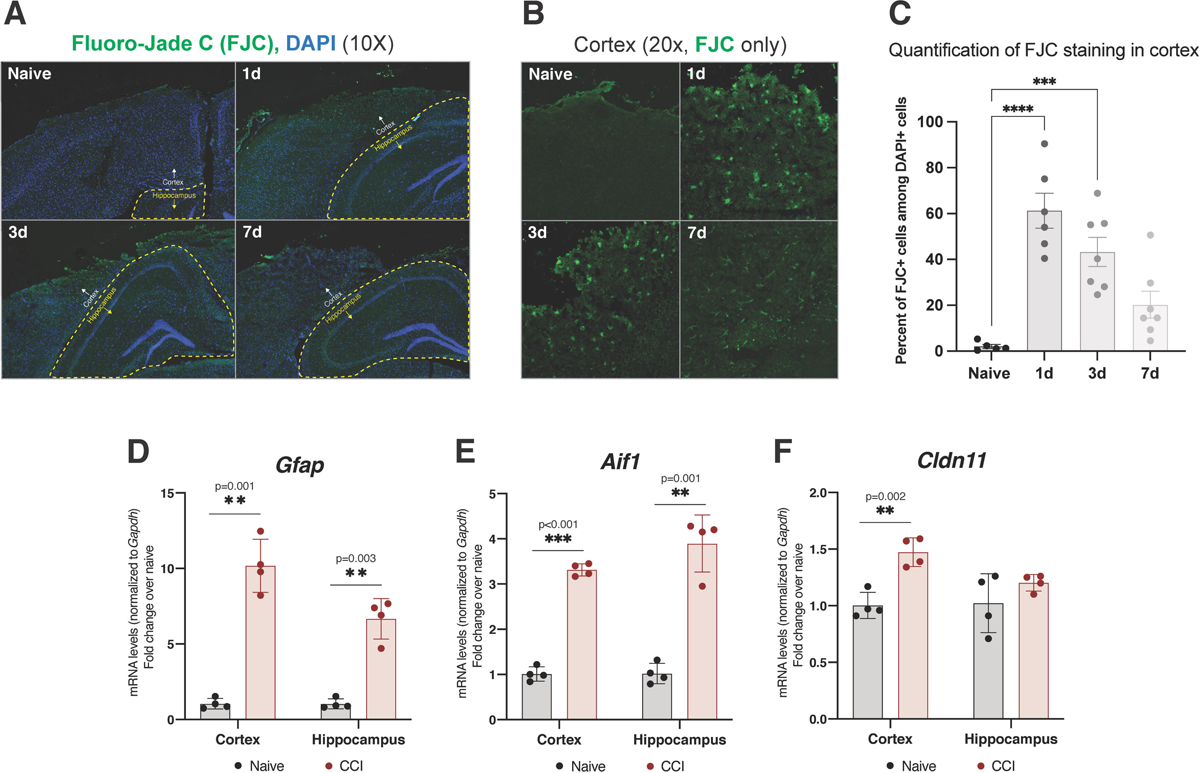
Regional specificity of cell death and glial proliferation after CCI. **(A)** Fluoro-Jade C (FJC) staining images at 10X magnification at multiple timepoints after CCI, representative of 3 separate mice (biological replicates) per group. The cortex and hippocampus are shown, with the hippocampus outlined in yellow. Degenerating neurons are visible as bright green puncta. **(B)** FJC staining images at 20X magnification in the ipsilateral cortex only. **(C)** Quantification of FJC staining in the cortex, represented as percentage of DAPI+ cells also positive for FJC signal. Statistical differences were determined by Ordinary one-way ANOVA followed by Bonferroni’s multiple comparisons test at an ⍺=0.05 significance level; ***, p<0.001; ****, p<0.0001. Error bars represent standard deviation among biological replicates. **(D)** *Gfap* gene expression via qPCR 3d after CCI, as an indicator of astrocytic proliferation. **(E)** *Aif1* gene expression (protein: IBA1) via qPCR 3d after CCI, as an indicator of microglial proliferation. **(F)** *Cldn11* gene expression (protein: oligodendrocyte-specific protein, OSP) via qPCR 3d after CCI, as an indicator of oligodendrocyte proliferation. For **D-F**, statistical comparisons are made to naïve (uninjured) tissues harvested and assayed alongside CCI tissues (two-tailed T-test; ⍺=0.05; **, p<0.01; ***, p<0.001). Each data point represents a separate mouse (biological replicate) and is representative of 3 technical replicates (qPCR wells). Error bars represent standard deviation among biological replicates.

### MAM activity is upregulated after brain injury

Our studies in AD models (patient fibroblasts and cellular/animal models) suggest that, through its ability to directly bind cholesterol, C99 accumulation in the ER induces the formation of MAM domains by stimulating the trafficking of cholesterol to the ER and the subsequent formation of lipid rafts (Pera et al 2017; Montesinos et al 2020c). Given the role of MAM in cellular lipid regulation, and previous reports of TBI stimulating lipid synthesis for membrane repair (Vance et al 2000; Adibhatla and Hatcher 2007), we hypothesized that brain injury could also induce increases in MAM-localized C99 to promote these processes. As previously published in the context of CCI (Mohamed et al 2021), these repair mechanisms appear to be triggered 3d after injury following resolution of initial edema. Indeed, 3d after CCI, we observed a marked increase in C99 levels in MAM fractions collected from both the ipsilateral cortex and hippocampus of injured mice relative to naïve controls (Fig. 2A, Fig. S1A). The occurred in the absence of observable alterations in the expression of β-secretase (gene *Bace1*), the enzyme that cleaves full-length APP to generate C99 (Fig. S1C).

**Figure 2:**
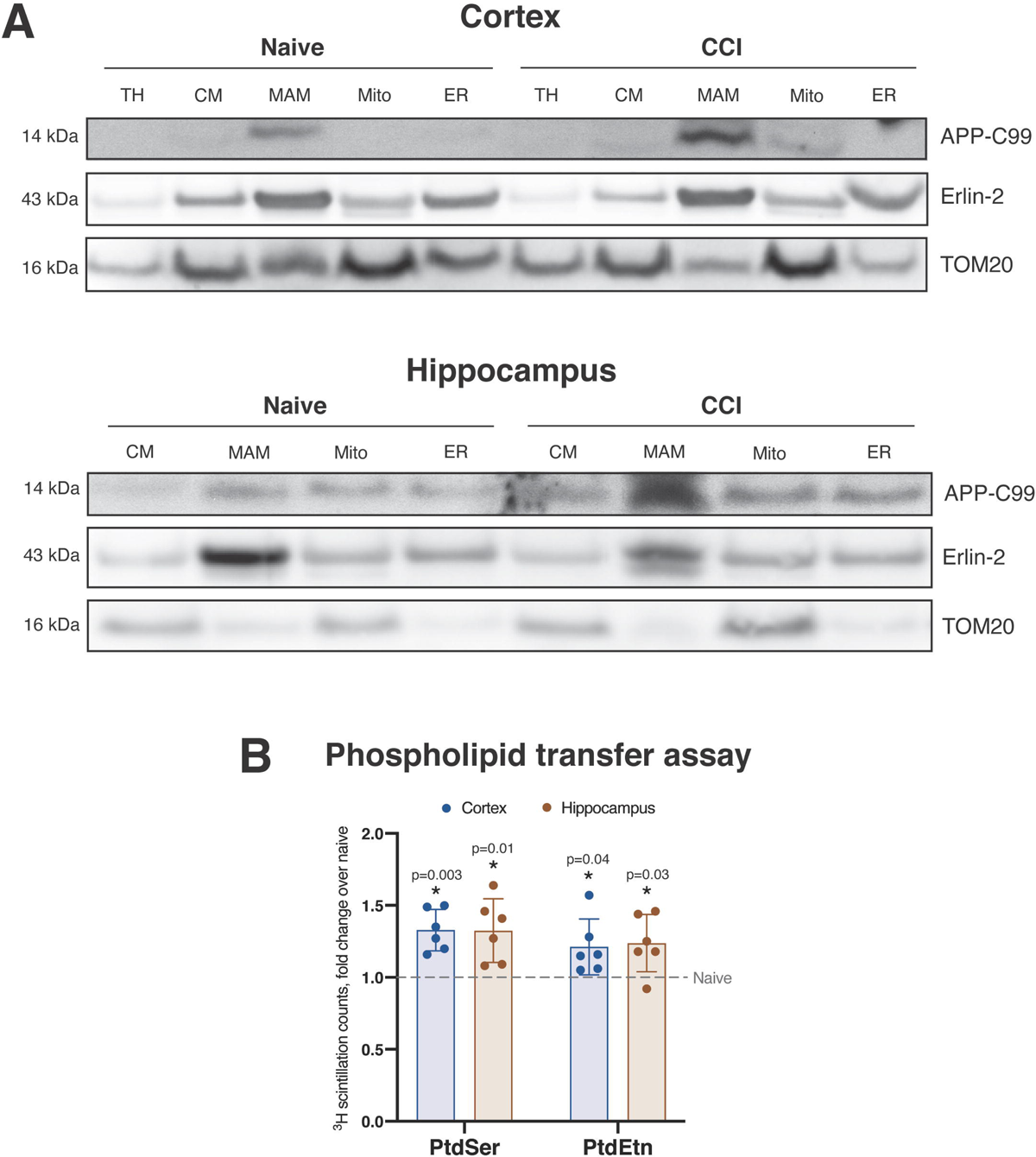
Increased localization of APP-C99 to MAM domains after CCI correlates with increased phospholipid transfer activity. **(A)** Subcellular fractionation of ipsilateral cortical and hippocampal homogenates 3d after CCI. CM, crude mitochondria fraction; MAM, mitochondria-associated ER membranes fraction; Mito, purified mitochondria; ER, bulk ER fraction. Erlin-2 and TOM20 serve as MAM and mitochondria markers, respectively. This western blot is representative of 3 biological replicates, each conducted with pooled tissues from 4 mice/group. **(B)** Analysis of phospholipid synthesis 3d after CCI. PtdSer, phosphatidylserine; PtdEtn, phosphatidylethanolamine. Each data point represents a separate mouse (biological replicate) and the average of 3 technical replicates. Statistical comparisons are made to naïve (uninjured) tissues from the same region assayed alongside CCI tissues (two-tailed T-test; ⍺=0.05; *, p<0.05). Error bars represent standard deviation among biological replicates.

We next sought to determine whether this result correlated with increases in MAM-specific enzymatic activities. We measured phospholipid synthesis and transfer between the ER and mitochondria, an established proxy measure of MAM activity (Vance 1990). Briefly, serine is incorporated into phosphatidylserine (PtdSer) by PtdSer synthase 1 (PSS1) at MAM domains, after which it is transferred to mitochondria for decarboxylation by PtdSer decarboxylase (PISD) to form phosphatidylethanolamine (PtdEtn) (Vance 2008). Upon pulsing crude mitochondria fractions with radiolabeled serine, we observed significantly increased production of radiolabeled PtdSer and PtdEtn in both the cortex and hippocampus of CCI mice compared to naïve controls, indicating an increase in MAM activity (Fig. 2B).

The formation of MAM induces the intracellular trafficking of cholesterol between membranes and the concomitant activation of sphingomyelinases (SMases) (Pera et al 2017), which hydrolyze sphingomyelin (SM) to ceramide, to facilitate this trafficking. To support the suggested increase in MAM activity after CCI, we measured SMase activity in hippocampal homogenates from CCI and naïve mice and observed a significant increase 3d after CCI (Fig. 3A). Remarkably, among the ≥5 mammalian SMases (Marchesini and Hannun 2004), only the expression of *Smpd5*, which encodes a putative mitochondria-associated neutral SMase (MA-nSMase) (Wu et al 2010), was significantly upregulated after TBI in both the cortex and hippocampus (Fig. 3B). In connection with its role of promoting cholesterol trafficking, SMase activity specifically correlates with cholesterol esterification by acyl-CoA:cholesterol acyltransferase 1 (ACAT1; gene *Soat1*) at MAM domains (Pera et al 2017). ACAT1 activity converts free cholesterol (FC) to cholesteryl esters (CEs) for storage in lipid droplets (LDs) (Area-Gomez et al 2012). In agreement with an increase in MAM activation after brain injury, we observed increased LDs in both the cortex and hippocampus after CCI compared to naïve controls (Fig. 3C). We confirmed these results by staining naïve and CCI tissues with Oil Red O, which identifies neutral lipid deposits (Mehlem et al 2013), and observed increased staining in both brain regions after injury (Fig. 3D). To validate this result at higher resolution and specificity, we imaged the vibrational energy of lipid-specific C-H bonds through Stimulated Raman Scattering (SRS) imaging 7d after injury (Shi et al 2018). This approach revealed a clustered signal reminiscent of granular lipid deposition, a pattern that was absent in naïve tissues (Fig. S2). Together, these results speak to the stimulation of MAM activity by brain injury, characterized by phospholipid synthesis, SMase activation and cholesterol esterification.

**Figure 3:**
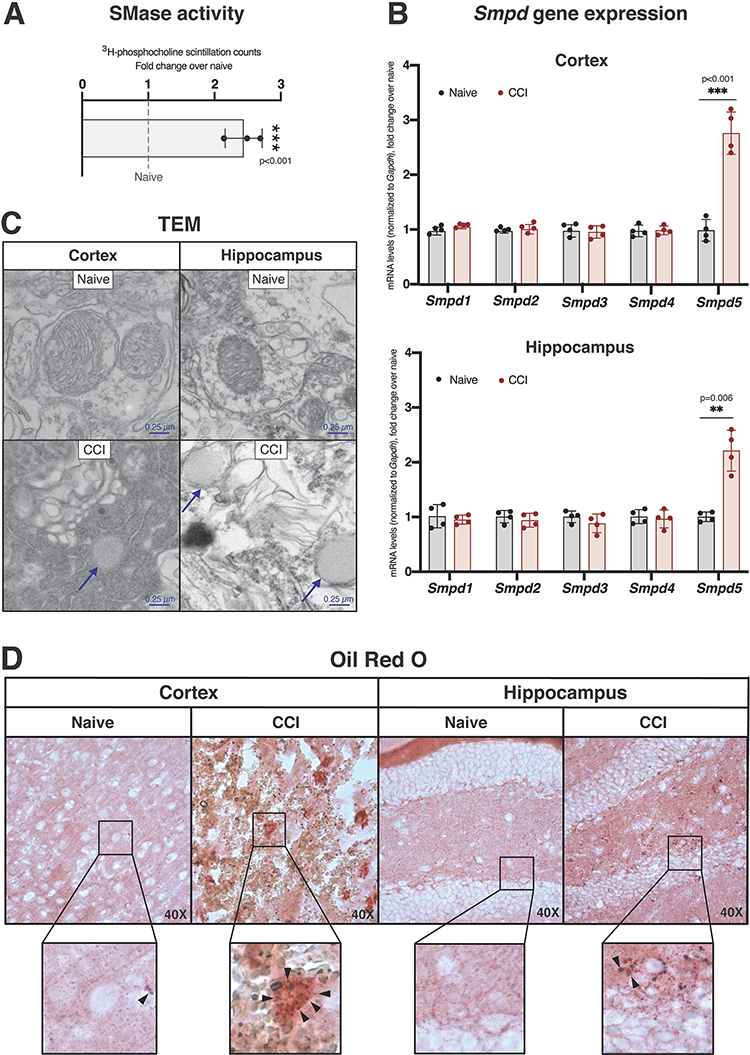
Cholesterol trafficking is upregulated after CCI in both the cortex and hippocampus. **(A)** Sphingomyelinase (SMase) activity assay in the hippocampus 3d after injury. Each data point represents a separate mouse (biological replicate) and the average of 3 technical replicates. **(B)** Gene expression analysis via qPCR of the *Smpd* genes, which encode the five mammalian SMase isoforms, 3d after CCI, normalized to *Gapdh*. Each data point represents a separate mouse (biological replicate) and the average of 3 technical replicates (qPCR wells). For **A** and **B**, statistical comparisons are made to naïve (uninjured) tissues from the same region assayed alongside CCI tissues (two-tailed T-test; ⍺=0.05; *, p<0.05; **, p<0.01). Error bars represent standard deviation among biological replicates. (**C**) Electron micro-scopy image 3d after CCI with visible lipid droplets, representative of 3 separate mice (biological replicates) per group. **(D)** Oil Red O staining of the ipsilateral cortex and hippocampus 3d after CCI, representative of 3 separate mice (biological replicates) per group. Lipid droplets are visible as bright red puncta.

### Alterations in the lipid composition of brain tissues coincide with the activation of MAM

MAM domains are a predominant hub for lipid regulation in the cell, as multiple lipid metabolic pathways converge at this single cellular locus (Vance 2014). Thus, increased MAM formation causes the activities of MAM-resident metabolic enzymes to be upregulated, manifesting as global alterations in the lipid composition of cellular membranes. As such, states of MAM upregulation and activation of relevant pathologies have become associated with specific lipidomic signatures. We thus conducted lipidomics analysis in our CCI model to ascertain progressive MAM-driven alterations in the cellular lipidome 1, 3 and 7 days 1, 3, and 7 days after injury (Fig. 4A). Consistent with the known correlation of MAM activity with cholesterol esterification and deposition of CE-rich LDs (Area-Gomez et al 2012), we observed significant increases in CE levels in all lipidomics data sets collected (Fig. 4A). This was consistent with elevated ratios of CE:FC (Fig. 4A), a recognized measure of inter-membrane cholesterol trafficking (Montesinos et al 2020c). These alterations were sustained in the cortex over the week-long period following injury while nearly normalizing in the hippocampus over the same time period.

**Figure 4:**
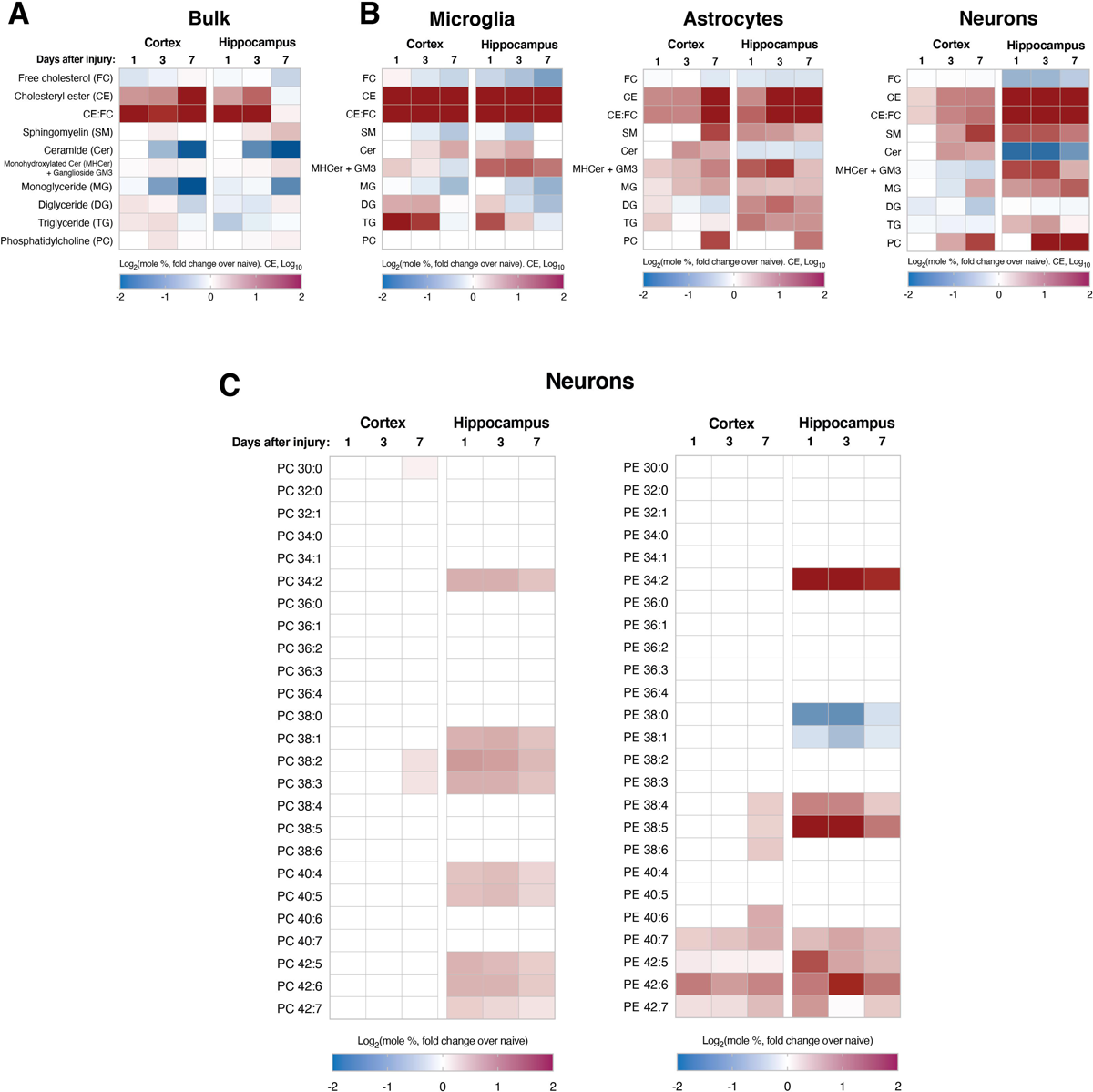
Lipidomic alterations in various lipid classes are observable in both bulk homogenates and purified cell type-specific populations after CCI. **(A, B)** Heatmap of significant differences in indicated lipid species as determined by lipidomics analysis at 1, 3 and 7 days after CCI (fold change over naïve), in (A) bulk cortical and hippocampal homogenates (n = 3 biological replicates) and (B) purified populations of microglia, astrocytes and neurons. **(C)** Lipidomics analysis of PtdCho (PC) and PtdEtn (PE) species in neurons purified from the ipsilateral cortex and hippo-campus after CCI. For **A-C**, only statistically significant differences between each time-point and naïve samples from the same region and/or cell type are shown (two-tailed T-test; ⍺=0.05; p<0.05).

To generate additional insight into the molecular origins of the observed lipidomic alterations, we assayed the expression of various regulators of cellular lipid uptake, trafficking and metabolism. The expression of *Abca1*, a lipid efflux transporter (Kim et al 2008), was increased significantly in both the cortex and hippocampus (Fig. S4A). We also observed increased expression of scavenger receptors and lipoprotein receptors. Specifically, *Cd36*, a scavenger receptor for free fatty acids (FAs) (Goldberg et al 2009), was robustly increased in both the cortex and hippocampus (Fig. S4C), and *Scarb1,* another scavenger receptor (Grampp et al 2017), was also increased in the cortex with a trend toward an increase in the hippocampus (Fig. S4C). Furthermore, the expression levels of *Ldlr* and *Lrp1*, lipoprotein receptors that transport cholesterol-rich lipoproteins (Foley and Esko 2010), were increased in the cortex but not in the hippocampus (Fig. S4B). Finally, we observed increased cortical expression (with a trend toward an increase in the hippocampus) of *Npc2*, encoding Niemann-Pick type C intracellular cholesterol transporter 2, a regulator of the inter-membrane transport of internalized cholesterol (Infante et al 2008) (Fig. S4B). This data validates our observations of stimulated cholesterol turnover following injury.

In addition to cholesterol species, CCI tissues also displayed significant increases in diglyceride (DG) and triglyceride (TG) levels (Fig. 4A/B). Elevations in these lipid species suggest a robust activation of lipid remodeling pathways upon injury. Specifically, we observed a notable increase in DG/TG species containing 16/18-carbon-long saturated and monounsaturated fatty acyl chains (e.g., DG 36:0, TG 50:0, etc.), generally associated with the activation of lipogenic pathways in the cell (Ameer et al 2014). Indeed, the expression of *Srebf1* [encoding sterol regulatory element-binding protein 1, SREBP1, a master regulator of sterol and fatty acid synthesis (Horton et al 2002)], *Fasn* (encoding fatty acid synthase, FAS) and *Acaca* (encoding acetyl co-A carboxylase-1, ACC1) were significantly increased in the cortex with a trend toward an increase in the hippocampus (Fig. S4D). These findings further support a substantial lipid metabolic response to brain injury, presumably to promote membrane repair.

The above findings were collected in bulk tissue homogenates. However, it is well known that the different cell types comprising the brain have vastly different metabolic profiles and carry out distinct functions during the response to injury. To determine the contribution of different cell types to the lipidomic alterations described above, we also characterized microglia, astrocytes and neurons isolated from both the cortex and hippocampus (Fig. S3) by lipidomics (Fig. 4B). Among the purified cell type-specific populations, microglia from both brain regions showed a robust remodeling of their lipid composition and indications of increased MAM formation. Specifically, microglia displayed consistent elevations in total CE levels (Fig. 4B) and multiple CE species (Fig. S4F), as well as higher CE:FC ratios (Fig. 4B), relative to control. Elevations in numerous CE species were also observable in astrocytes and neurons, especially at 7d, and were more apparent in the hippocampus than in the cortex (Fig. S4F). In agreement with these cholesterol alterations, microglia from CCI tissues also displayed significant reductions in SM levels with concomitant increases in ceramide (Fig. 4B), indicative of upregulated SMase activity. Microglia also displayed a marked decrease in monoglyceride (MG) levels (Fig. 4B), consistent with the known association between inflammation and monoglyceride lipase (gene *Mgll*) activation (Habib et al 2019). The increased expression of FA scavenger receptor *Cd36*, as described above, in both the cortex and hippocampus (Fig. S4C) also supports the aforementioned indications of microglial proliferation owing to the known role of CD36 in microglial proliferation (Helming et al 2009).

When DGs and TGs were analyzed at the cell type-specific level in order to better understand the observed bulk increases, it was observed that these species were most significantly elevated in cortical microglia and hippocampal astrocytes (Fig. 4B), potentially to support astroglial and microglial proliferation. Furthermore, levels of phosphatidylcholine (PtdCho), the most abundant phospholipid species in the brain (Vance 2014), were increased significantly in astrocytes and neurons. Specifically, neurons from both brain regions displayed specific increases in PtdCho and PtdEtn species containing long polyunsaturated fatty acyl chains, especially at 1d and 3d (Fig. 4C). This suggests an increase in the desaturation index of neuronal membranes after injury, most robustly observable in the hippocampus. Altogether, the cell type-specific lipidomics data helps elucidate the bulk assay results described above and provides specific evidence for disrupted homeostasis of each of the assayed neural cell types.

### Mitochondrial respiration is not significantly affected after injury

We have previously reported that MAM upregulation results in mitochondrial dysfunction in AD models (Pera et al 2017). Similarly, mitochondrial dysfunction is a widely-reported phenotype in experimental TBI (Cheng et al 2012; Hiebert et al 2015). We therefore sought to determine the time-course and regional specificity of respiratory alterations in our TBI model by Seahorse microplate respirometry (Agrawal et al 2020). Surprisingly, we did not observe any significant alterations in oxygen consumption rate in mitochondria isolated from CCI brains at 1, 3, or 7 days after injury in either the cortex or hippocampus (Fig. 5A/B). This was observable when assaying both complex I (CI)-mediated respiration (indicative of pyruvate oxidation, measured in the presence of pyruvate with malate as a complex II inhibitor, Fig. 5A) and complex II (CII)-mediated respiration (indicative of fatty acid oxidation, FAO, measured in the presence of CII substrate, succinate, with rotenone as a CI inhibitor, Fig. 5B) (Agrawal et al 2020). We confirmed this through histological analysis of succinate dehydrogenase (SDH, full name of CII) and cytochrome *c* oxidase (COX, complex IV) activities (Ross 2011), both of which showed no changes in staining intensity in either region at all timepoints tested (Fig. 5C/D and Fig. S5A/B). We also measured the expression of the OxPhos respiratory complexes in both brain regions at these timepoints and, as with the enzymology, did not observe significantly consistent alterations in expression upon injury (Fig. S5F/G).

**Figure 5:**
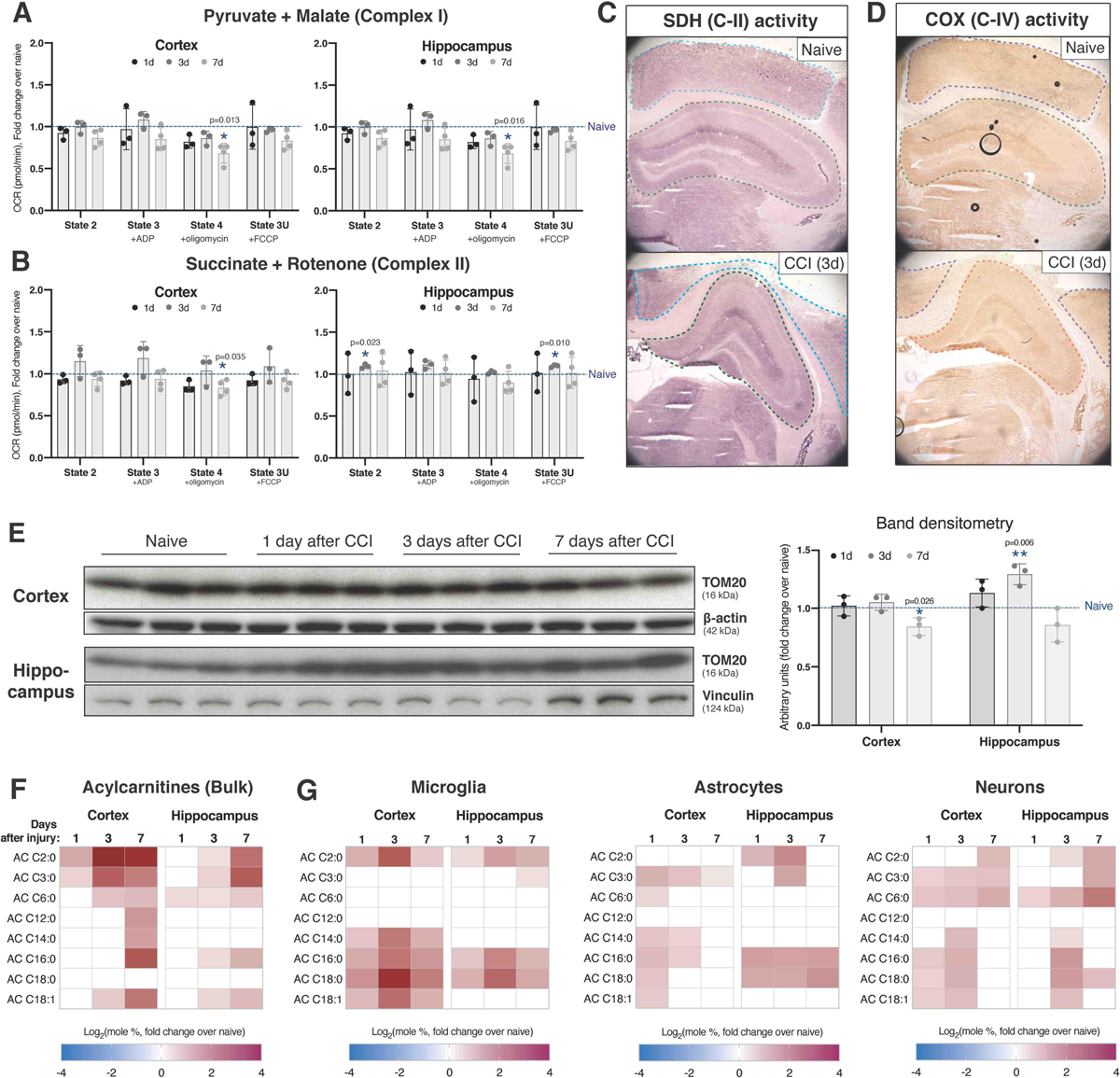
Cortical and hippocampal mitochondria display a moderate switch in substrate preference without impairments in bioenergetics. **(A,B)** Mitochondrial oxygen consumption rate (OCR) measured in the Seahorse XF Analyzer in crude mitochondrial fractions collected at the indicated time-points from the indicated ipsilateral brain regions. OCR values are shown by respiratory state: State 2, baseline; State 3, after ADP addition to stimulate electron flow; State 4, after oligomycin (ATP synthase inhibitor) addition to determine the fraction of oxygen consumption attributable to ATP production vs. proton leak; State 3-uncoupled (3U), after trifluoromethoxy carbonylcyanide phenylhydrazone (FCCP) addition in order to uncouple electron flow and oxygen consumption from ATP production, to determine maximal respiration levels. (A) Respiration in the presence of pyruvate and malate (complex-II inhibitor) in order to assay the contribution of complex-I (i.e., pyruvate oxidation) to cellular respiration. (B) Respiration in the presence of succinate (complex-II substrate) and rotenone (complex-I inhibitor) in order to assay the contribution of complex-II (i.e., fatty acid oxidation) to cellular respiration. Each data point represents a separate biological replicate (pooled tissues from 4 mice) and the average of 5 technical replicates (assay plate wells). **(C)** Succinate dehydrogenase (SDH, complex-II) activity staining images in the ipsilateral cortex (outlined in blue) and ipsilateral hippocampus (outlined in yellow) 3d after CCI, representative of 3 separate mice (biological replicates) per group. **(D)** Cytochrome C oxidase (COX) activity staining images in the ipsilateral cortex (outlined in blue) and ipsilateral hippocampus (outlined in yellow) 3d after CCI, representative of 3 separate mice (biological replicates) per group. **(E)** Expression of TOM20 (a marker for mitochondrial mass) in cortical and hippocampal homogenates at the indicated time-points after injury. Each lane is a separate mouse (biological replicate). β-actin is the loading control for the cortex and vinculin is the loading control for the hippocampus. Band densitometry is shown to the right. **(F)** Bulk lipidomics analysis of total acylcarnitines (ACs), which become elevated with increased fatty acid β-oxidation. This data is averaged from 3 biological replicates. **(G)** Lipidomics analysis of total ACs in astrocytes and microglia from both the cortex and hippocampus. For **A, B, E, F** and **G**, statistical comparisons are made to naïve (uninjured) tissues from the same region and/or cell type assayed alongside CCI tissues (two-tailed T-test; ⍺=0.05; *, p<0.05; **, p<0.01). For the lipidomics data in **F** and **G**, only statistically significant differences are indicated (p<0.05). Error bars represent standard deviation.

As a control for these assays, we measured various parameters related to mitochondrial mass. We first assayed the expression of TOM20 by western blot (Fig. 5E). Compared to naïve mice, we observed a slight but significant decrease in the cortex at 7d and an increase in the hippocampus at 3d. We also assayed the ratio of mitochondrial DNA (mtDNA) to nuclear DNA (nDNA) through qPCR analysis of *Cox1* (encoded by the mitochondrial genome) and *Gapdh* (encoded by nuclear genome) gene levels. Of note, the ratio was unchanged in the cortex but was significantly decreased in the hippocampus at 3d and 7d (Fig. S5C). Similarly, the expression of *Ppargc1a*, encoding PGC-1⍺, a regulator of mitochondrial biogenesis (Jornayvaz and Shulman 2010), was significantly decreased in both brain regions at 3d (Fig. S5D). TBI also has reported associations with hypoxic stress and a consequent reduction in pyruvate oxidation by mitochondria (Maloney-Wilensky et al 2009; Fuhrmann et al 2019). In agreement with this, the expression of *Hif1a,* encoding the transcription factor hypoxia-inducible factor 1-⍺ (HIF1⍺), was elevated in the cortex but not in the hippocampus (Fig. S6A). A similar result was observed for HIF target, *Pdk1*, encoding pyruvate dehydrogenase kinase 1 (PDK1) (Fig. S6B). PDK1 inhibits pyruvate dehydrogenase (PDH) activity (Jha et al 2012) and thus reduces the use of pyruvate as a mitochondrial fuel source, which can cause mitochondria to begin oxidizing alternative fuel sources, such as FAs. In support of this possibility, acylcarnitines (ACs), intermediate metabolites of mitochondrial FAO (Longo et al 2016), showed significant increases in both the cortex and hippocampus at multiple timepoints (Fig. 5F). However, the expression of *Cpt1a*, which encodes carnitine palmitoyltransferase 1⍺ (CPT-1⍺), the enzyme that enables transport of fatty acyl-coAs into the mitochondrial matrix for oxidation (McGarry and Brown 1997), was significantly increased only in the cortex (Fig. S5E). When lipidomics analysis of ACs was repeated in purified cell type-specific populations, we observed that while microglia and astrocytes were certainly contributing to the bulk trends, cortical and hippocampal neurons displayed the most robust increases in these AC species (Fig. 5G).

Altogether, our results suggest that elevations in MAM-localized C99 after brain injury are associated with increased MAM activity. This correlates with subsequent changes in the regulation of cellular lipid metabolism and the composition of cellular membranes, both at the bulk and cell type-specific levels. Nonetheless, mitochondrial respiratory capacity in minimally modulated.

## Discussion

The epidemiological connection between TBI and AD has been described in numerous studies, but the molecular connection has not been completely determined. Our group has previously reported that AD models display upregulated function of MAM domains (Area-Gomez et al 2012) at early disease stages. In this work, we present data suggesting that MAM functions are also upregulated in a mouse model of TBI. This upregulation correlates with significant modulations in lipid metabolism, observable at both the bulk and cell type-specific levels, without marked impairments in mitochondrial respiration.

Alterations in APP processing, a cardinal feature of AD, have been examined in multiple TBI studies. Deposition of Aβ occurs in humans after TBI (Smith et al 2003), albeit in a minority of patient samples in some studies (Ikonomovic et al 2004). This suggests that other APP processing products may be involved in the development of AD phenotypes after injury. Indeed, we have demonstrated that AD tissues and cell models display pathogenic accumulation of Aβ precursor, C99 (Pera et al 2017), and that C99 elevations are sufficient to upregulate MAM functionality (Montesinos et al 2020c). This finding has has been reported in TBI studies as well (Chen et al 2004; Cartagena et al 2016) and has been attributed to elevated expression of APP (C99 precursor) and BACE1 (the enzyme that cleaves APP to form C99) in damaged brain areas (Ciallella et al 2002; Blasko et al 2004; Chen et al 2004). C99 has also been demonstrated to correlate more strongly with brain damage than Aβ does (Rockenstein et al 2005; Jiang et al 2010; Tamayev et al 2012; Lauritzen et al 2012), suggesting a critical and potentially early role of C99 in TBI pathogenesis. In this work, we show that elevations of C99 are most observable in MAM domains of the ER 3d after a single CCI injury. This timing is consistent with reports from other groups showing that, compared to sham animals, TBI animals display significant structural alterations in white matter beginning at 3d and partially resolving by 7d, without any overt behavioral changes (Mohamed et al 2021). Thus, C99 elevations are among the molecular pathologies that are triggered in the early phase following brain injury.

The formation of MAM domains in the ER generates localized “signaling” platforms where a specific subset of lipid metabolic enzymes is recruited and activated to adapt the cellular lipidome to environmental changes. Activation of cholesterol turnover pathways is an especially key adaptation that prevents cellular cholesterol levels from surpassing a toxic threshold. The enzymes responsible for this homeostatic regulation are acyl-CoA cholesterol acyltransferase 1 (ACAT1, gene *SOAT1*), which enables removal of excess cholesterol from membranes by generating cholesteryl esters (CEs) (Chang et al 2009), and ER lipid raft-associated protein 2 (ERLIN2/SPFH2, gene *ERLN2*), which inhibits the *de novo* synthesis of additional cholesterol by inducing degradation of 3-hydroxy-3-methylglutaryl-CoA reductase (HMGCR, gene *HMGCR*) (Jo et al 2011). Consistent with this role of MAM in detoxifying excess cellular cholesterol and previous reports of TBI triggering cholesterol accumulation (Cartagena et al 2008), our lipidomics analysis of TBI tissues showed progressive reductions in non-esterified cholesterol and concomitant increases in cholesteryl esters (CEs), suggesting increased mobilization of cholesterol from cellular membranes, similar to AD (Pera et al 2017; Montesinos et al 2020c). Our findings are also consistent with previously reported increases in the expression of genes involved in cholesterol mobilization, such as cholesterol transporter APOE (de Chaves et al 1997; Cartagena et al 2008; Tweedie et al 2016), lipid transporter LDLR (de Chaves et al 1997), and lipid efflux pump ABCA1 (Cartagena et al 2008; Loane et al 2011; Jasmin et al 2014). Indeed, the ε4 allele of ApoE (ApoE4) – the strongest genetic risk factor for AD – has been shown in some studies to perpetuate pathogenic cascades following TBI (McFadyen et al 2021). Thus, cholesterol metabolism pathways, including the specific steps carried out by MAM-resident enzymes, are likely intimately involved in TBI pathogenesis, similar to AD.

In addition to the bulk trends described above, cholesterol alterations were also observable in purified cell type-specific populations. Lipidomics analysis in neurons, astrocytes and microglia collected from TBI tissues showed that increases in CEs were particularly prominent in microglia. Our findings are consistent with early reports of lipid accumulation in nerve injury models, where lipid droplets (LDs) were observed in brain macrophages to presumably support lipid trafficking for axon regeneration (Boyles et al 1989; Goodrum et al 1994). AD-associated pro-inflammatory microglia display cholesterol accumulation as well (Feringa and van der Kant 2021), and decreased lipid clearance in astrocytes and microglia from the AD-associated *APOE4* background has also been described (Tcw et al 2022). At the molecular level, cholesterol molecules in membranes are closely bound to sphingomyelin (SM), an important cellular sphingolipid (Slotte 1999). Indeed, cholesterol and sphingolipid metabolic pathways are co-regulated by shared master regulators (Gulati et al 2010). It thus follows that, similar to cholesterol alterations, sphingolipid alterations detectable by lipidomics were most prominent in microglia relative to other cell types in our TBI model. In agreement with TBI lipidomics studies by other groups (Roux et al 2016; Barbacci et al 2017), we observed reduced SM levels and increased ceramides, suggesting increased hydrolysis of SM and conversion to ceramide by cellular sphingomyelinases (SMases). We generated support for this possibility in our model by observing elevations in SMase enzymatic activity (Fig. 3A) and *Smpd5* gene expression (Fig. 3B). Interestingly, *Smpd5* is predominantly expressed in microglia relative to astrocytes and neurons (Zhang et al 2014) and, in our model, only microglia displayed the SM and ceramide alterations that are consistent with SMase activation (Fig. 4B). Of note, multiple activators of SMase activity [such as tumor necrosis factor-⍺ (Chatterjee 1994), arachidonic acid (Jayadev et al 1994), and glutamate (Novgorodov et al 2018)] are established components of TBI pathology. Furthermore, it is well known that microglial activation and the regulation of subsequent inflammatory cascades are closely modulated by changes in microglial lipid composition (Ruysschaert and Lonez 2015). For example, ligand-mediated activation of toll-like receptors results in the transient upregulation of SMases (Köberlin et al 2015) and the stimulation of cholesterol trafficking and esterification (Wong et al 2020). Indeed, pro-inflammatory microglia display increases in ceramides, CEs and LDs (Marschallinger et al 2020). In light of this association between cholesterol and sphingolipid alterations and microglial activation, and the wealth of evidence demonstrating a major contribution of neuroinflammatory pathways to AD pathogenesis, the upregulation of SMase activity (and cholesterol esterification) after TBI episodes represents a point of convergence between TBI and AD pathogenic mechanisms (Kinney et al 2018; Loving and Bruce 2020). Of note, a human homolog of *Smpd5* has not been characterized; thus, the relevance of these findings to human TBI requires further investigation.

The activation of glial cells is also associated with the accumulation of triglycerides (TGs) (Feingold et al 2012), which was observable at the bulk level in this study, especially at 1d and 3d after injury (Fig. 4A). In cell type-specific analyses, TG increases were most prominent in microglia at 1d and 3d, but more sustained (until 7d) in hippocampal astrocytes (Fig. 4B). Pro-inflammatory macrophages are known to increase the synthesis of fatty acids (FAs) and TGs and reduce FA oxidation, causing LDs and long-chain acylcarnitines (ACs) to accumulate (Castoldi et al 2020). Anti-inflammatory states are associated with progressively reduced TG synthesis and increased FA consumption, resulting in the accumulation of short-chain ACs over time (Koves et al 2008; Yang et al 2021). In agreement with these known trends, microglia in our CCI model displayed significant increases in long-chain AC levels that progressively diminished by 7d (Fig. 5G). Concomitantly, we observed progressive increases short-chain ACs over time (Fig. 5G), suggesting an eventual metabolic shift toward FA oxidation, as associated with anti-inflammatory phenotypes. Interestingly, when we measured bulk gene expression of lipid efflux pumps *Abca1* [enriched in astrocytes (Zhang et al 2014)] and *Abcg1* [enriched in microglia (Zhang et al 2014)], only *Abca1* was elevated after TBI (Fig. S5A). The same finding has been reported by other groups in models of neural injury as well (Cartagena et al 2008; Jasmin et al 2014). Altogether, these results suggest that TG phenotypes during the acute phase after injury derive from both microglial and astrocytic responses.

When the lipidomics results in post-CCI neuronal populations were analyzed, the most marked alterations were in phospholipids. Specifically, we observed progressive increases in PtdCho and PtdEtn molecules containing polyunsaturated fatty acids (PUFAs), which increase membrane fluidity (Yang et al 2011). This was observable throughout the week-long period following injury in only the hippocampus for PtdCho and in both the cortex and hippocampus for PtdEtn. These findings support previous reports of PtdCho and PtdEtn alterations in TBI models and the hypothesized connection to cellular membrane disruption after injury (Pasvogel et al 2008; Ojo et al 2019). Increases in phospholipid desaturation are induced by the activation of specific acyl-coA synthases (ACSLs), such as ACSL4 and ACSL6, which preferentially activate PUFAs for incorporation into phospholipids (Chouinard-Watkins and Bazinet 2018; Kuwata and Hara 2019). Of note, ACSLs are MAM-resident enzymes (Lewin et al 2002). Thus, it is possible that increased MAM activity is at the heart of potentially increased ACSL4/6 activity. The phospholipid alterations described here represent another point of convergence with AD pathogenesis, as a number of sporadic AD (SAD)-associated genetic polymorphisms are in genes regulating phospholipid transport and metabolism (Hardy 2017). Therefore, connections between the early stages of TBI and AD pathogenic mechanisms can be drawn from alterations in multiple lipid classes. Of note, regarding the brain regional specificity of these lipidomic alterations, we expected the cortex to display the greatest changes due to the physical disruption of this brain region in the CCI model employed here. While this was indeed the case for some of the studied lipid classes, the hippocampus displayed greater disturbances in CEs and PUFA-containing phospholipids. The origins of such regional specificity in lipid metabolic alterations following brain injury merit further investigation.

The final dataset presented in this study centers around mitochondrial bioenergetic homeostasis after CCI. Despite previous reports discussing mitochondrial bioenergetic impairments after TBI (Hiebert et al 2015), we did not observe significant impairments in mitochondrial respiration in our model (Fig. 5A/B). This discrepancy can likely be explained by methodological differences between our studies and other studies in the field. It merits mentioning that our data suggests a partial shift in OxPhos substrate preference away from pyruvate oxidation toward the oxidation of FAs (Fig. 5F/G). This is in agreement with previous reports (Kilbaugh et al 2015; Kilbaugh et al 2016). It is possible that this shift in fuel source represents a counterbalancing mechanism to prevent bioenergetic failure. Indeed, reduced mitochondrial pyruvate oxidation has been described as a cellular mechanism of preventing excitotoxicity (Divakaruni et al 2017). If sustained, this potential increased in mitochondrial FA oxidation could become detrimental for neuronal functionality (Schönfeld and Reiser 2013). Nonetheless, mitochondrial bioenergetics were not significantly disturbed within the injury parameters and assay window of this study.

In summary, we report here that a single CCI episode induces transient elevations in C99 and the formation and activation of MAM domains in the ER. We also report that lipidomic alterations after TBI are observable at both the bulk and cell type-specific levels, are especially prominent in microglia, and are consistent with the activation of MAM-specific lipid metabolic activities. It is likely that, after multiple injuries, dysregulation of these pathways would be sustained to the point where neuronal homeostasis is impaired, resulting in functional impairments. This would be a chronic phenotype, resembling that of AD. Our preliminary data lay the groundwork for an important role of MAM in lipid metabolic disturbances during the acute phase following brain injury, as well as for novel perspectives considering TBI as an environmental cause of AD.

## Supporting information

Lipidomics source data - astrocytes

Lipidomics source data - bulk

Lipidomics source data - microglia

Lipidomics source data - neurons

## Additional files

**Lipidomics source data.pdf (separate files for bulk, microglia, astrocytes and neurons)**: raw lipidomics data values (depicted as fold change over naïve samples) represented in the lipidomics heat maps in this manuscript.

## Statements and Declarations

### Author contributions

Conceived the project: EA-G. Designed experiments: EA-G and RRA. Generated data for most of the experiments: RRA. Provided CCI equipment and training: SGK. Critically assisted with Seahorse analysis: DL. Collected/analyzed lipidomics data: YX, TDY, RRA and EA-G. Conducted/analyzed SRS imaging studies: LS, DS, WM, RRA, RJD and EA-G. Conducted/analyzed Fluorojade-C staining: HZ and RRA. Conducted/analyzed COX/SDH staining: VE and RRA. Conducted/analyzed EM imaging: LGC, FPM, RRA, EA-G and RJD. Wrote the manuscript: RRA. Critically edited the manuscript: EA-G. Approved final version of the manuscript: all authors.

## Acknowledgements

We thank Kevin Velasco, Tzong-Shiue Yu and Yacine Tensaouti for laboratory assistance. We thank Renu Nandakumar for assistance with the lipidomics analysis, Sana Chintamen for advice on brain dissociation, Kristy Brown for assistance with preliminary electron microscopy experiments, and Patricia Washington and Stephanie Siegmund for assistance with preliminary Seahorse assays. Finally, we thank Kirstin Tamucci, Jorge Montesinos and Cristina Guardia-Laguarta for feedback on the manuscript, and all members of the Area-Gomez and Deckelbaum labs for helpful discussions.

## Data availability

Source data files have been submitted to the journal as requested. The lipidomics datasets supporting the conclusions of this article are included as a supplemental file. All reagents used in this work are commercially available, with manufacturer catalog numbers included in the Methods.

## Ethics approval

All animal husbandry was conducted in accordance with the Guide for the Care and Use of Laboratory Animals published by the National Institutes of Health. Specific procedures were approved by the Institutional Animal Care and Use Committee at Columbia University (protocols AC-AAAO5307 and AC-AAAY6450).

---

**Supplemental Figure 1:**
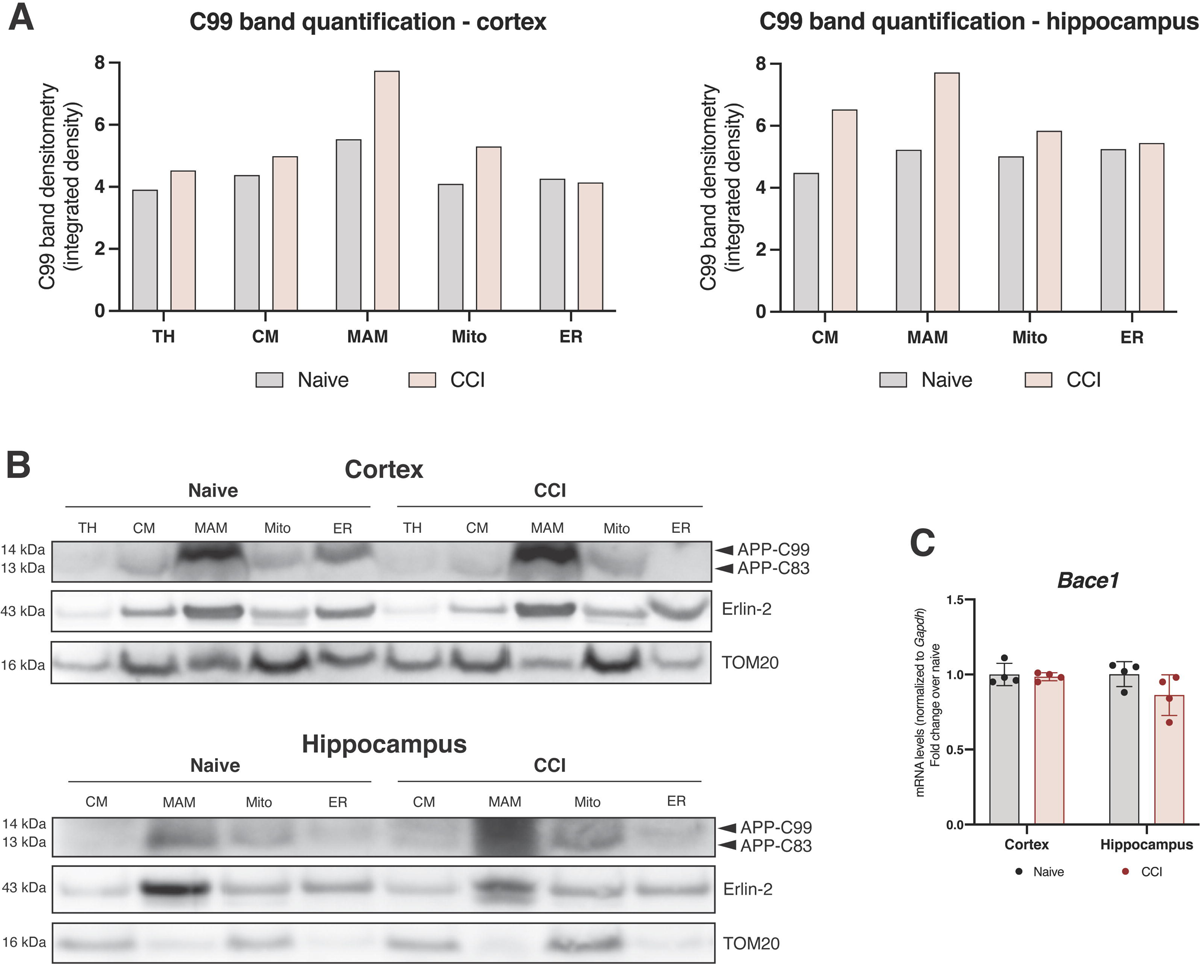
APP processing after CCI. **(A)** Quantification of C99 band intensity in both the cortex and hippocampus. **(B)** Subcellular fractionation of ipsilateral cortical and hippocampal homogenates 3d after CCI. TH, total homogenate; CM, crude mitochondria fraction; MAM, mitochondria-associated ER membranes fractions; Mito, purified mitochondria fraction; ER, bulk ER fraction. Erlin-2 and TOM20 serve as MAM and mitochondria markers, respectively. This western blot is representative of 3 biological replicates, each conducted with pooled tissues from 4 mice/group. **(C)** Gene expression analysis via qPCR of *Bace1*, which encodes β-secretase (the enzyme responsible for C99 production from full-length APP), 3d after CCI, normalized to *Gapdh*. Each data point represents a separate mouse (biological replicate) and the average of 3 technical replicates (qPCR wells). Error bars represent standard deviation among biological replicates. Statistical comparisons are made to naïve (uninjured) tissues from the same region harvested and assayed alongside CCI tissues (two-tailed T-test, ⍺=0.05).

**Supplemental Figure 2:**
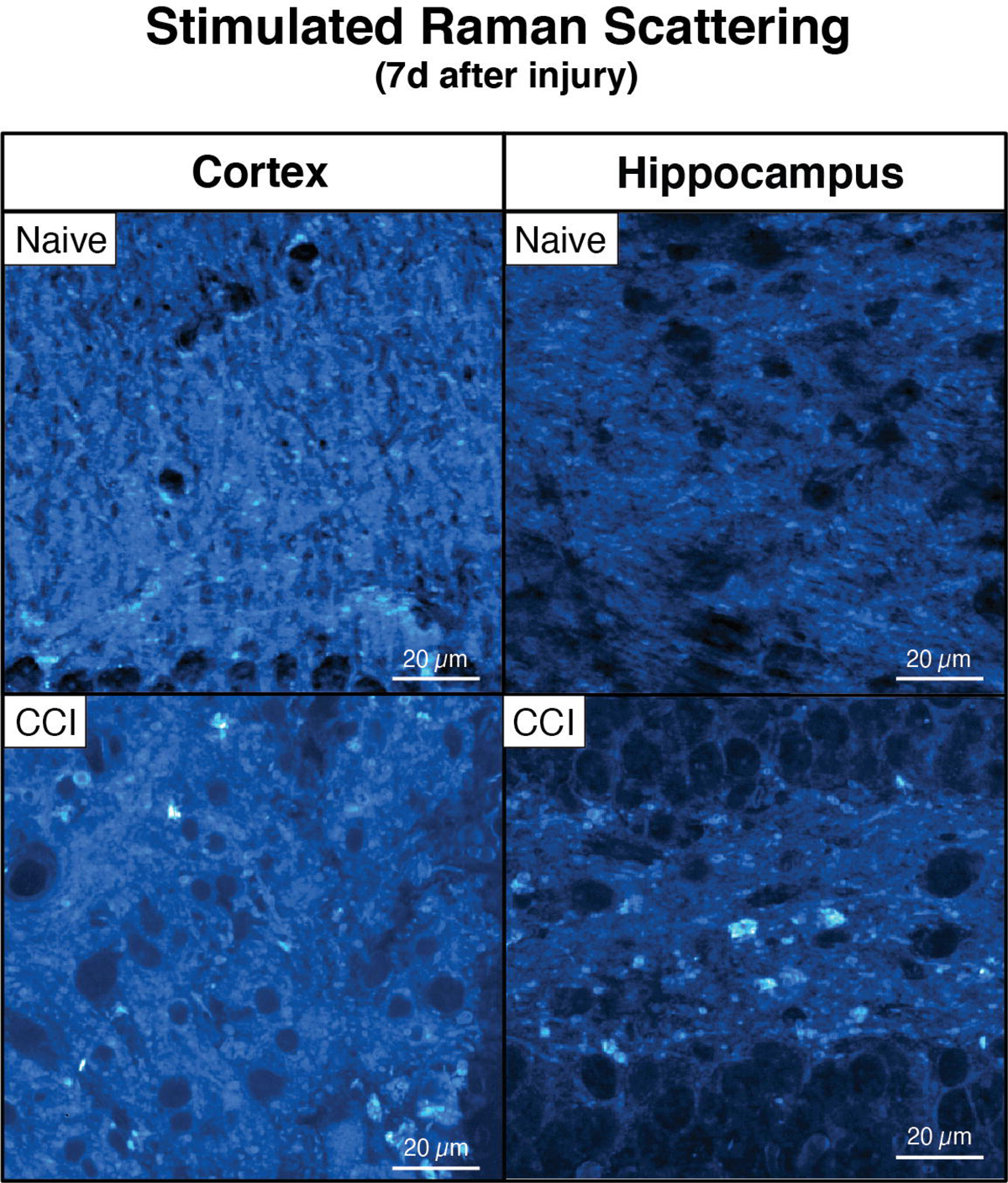
Stimulated Raman Scattering (SRS) imaging of lipid-specific C-H bonds 7 days after CCI. Stimulated Raman scattering (SRS) microscopy image of C-H bonds in the lipid channel 7d after CCI, representative of 3 separate mice (biological replicates) per group. Lipid clusters are visible as bright puncta.

**Supplemental Figure 3:**
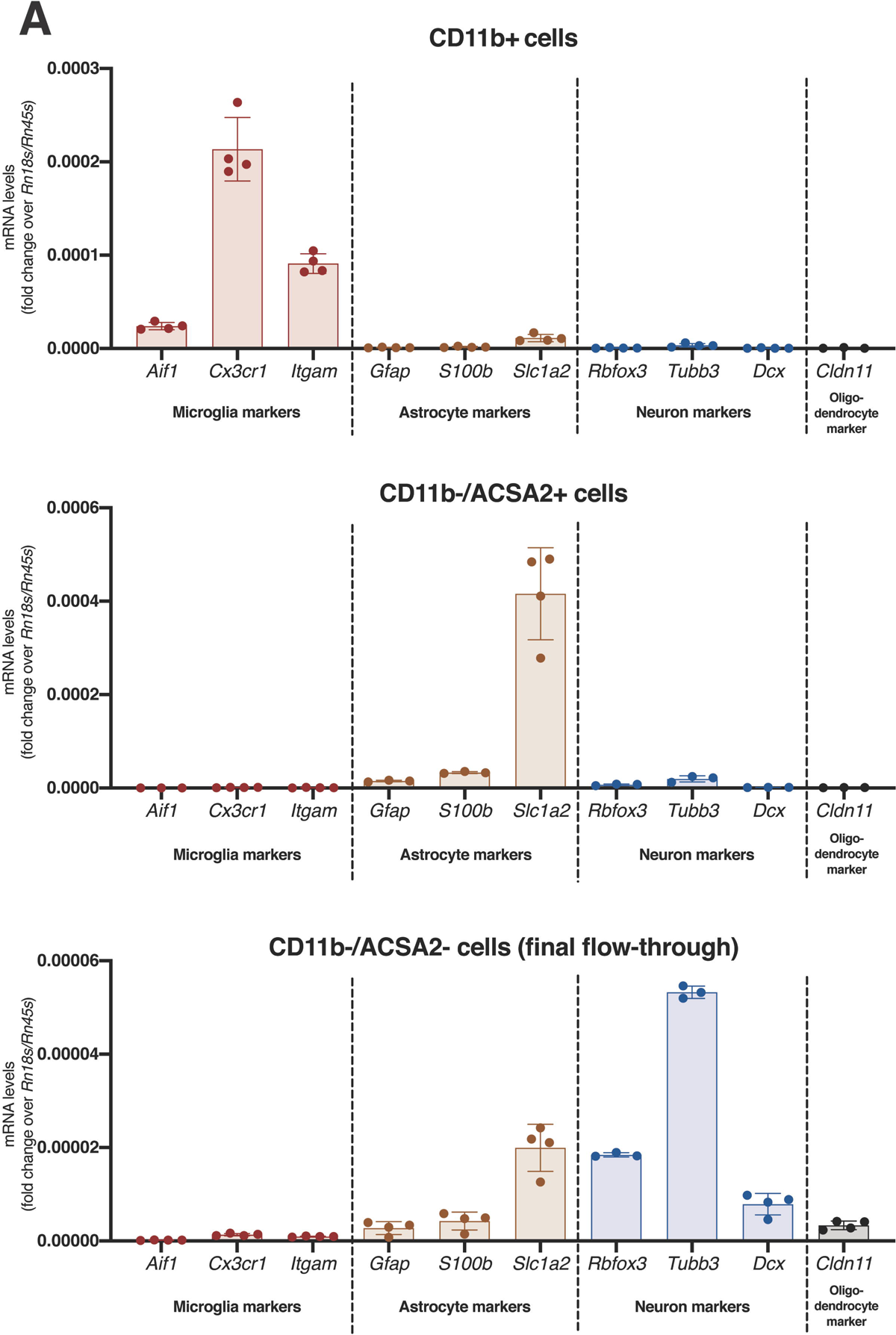
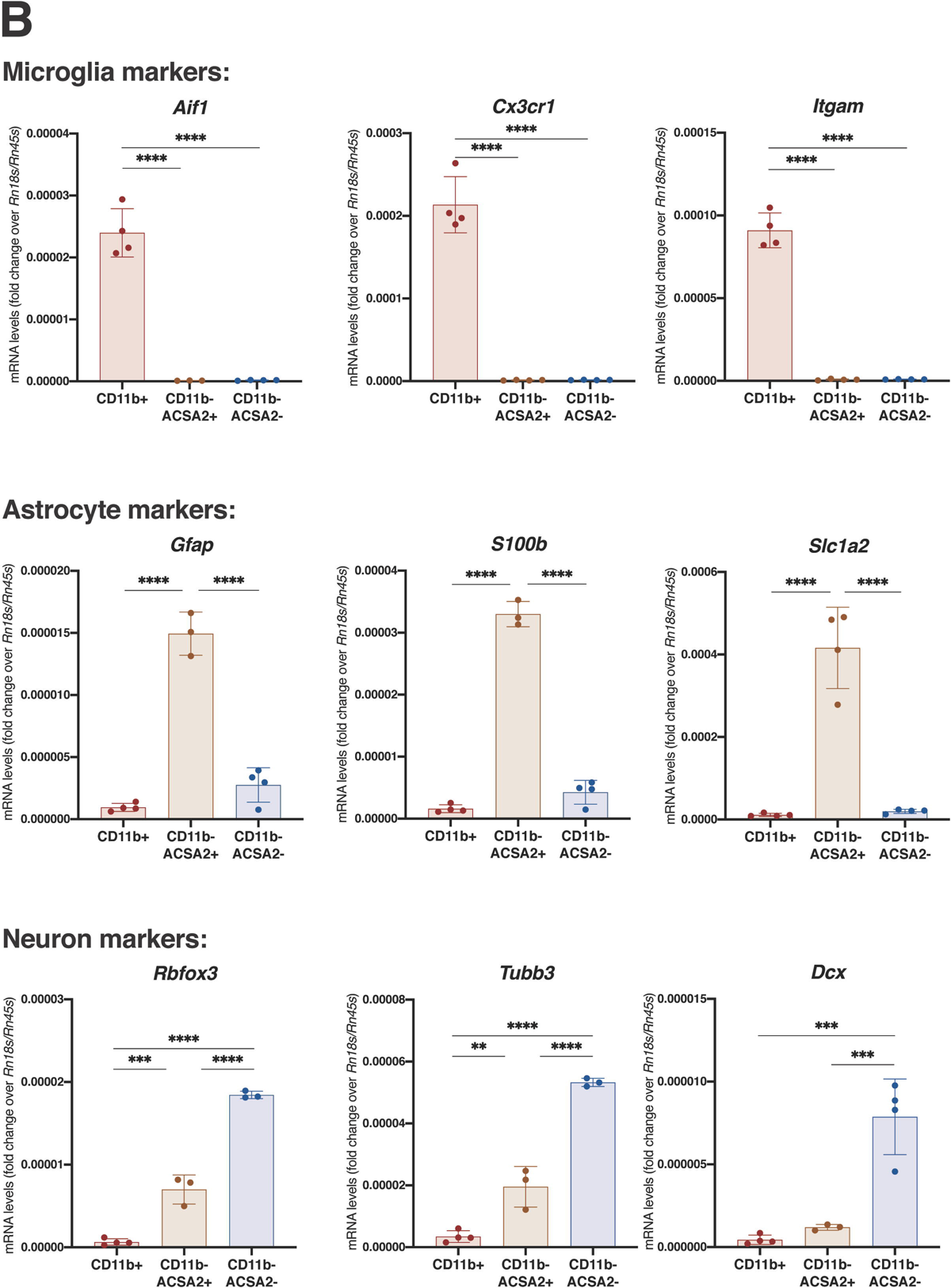
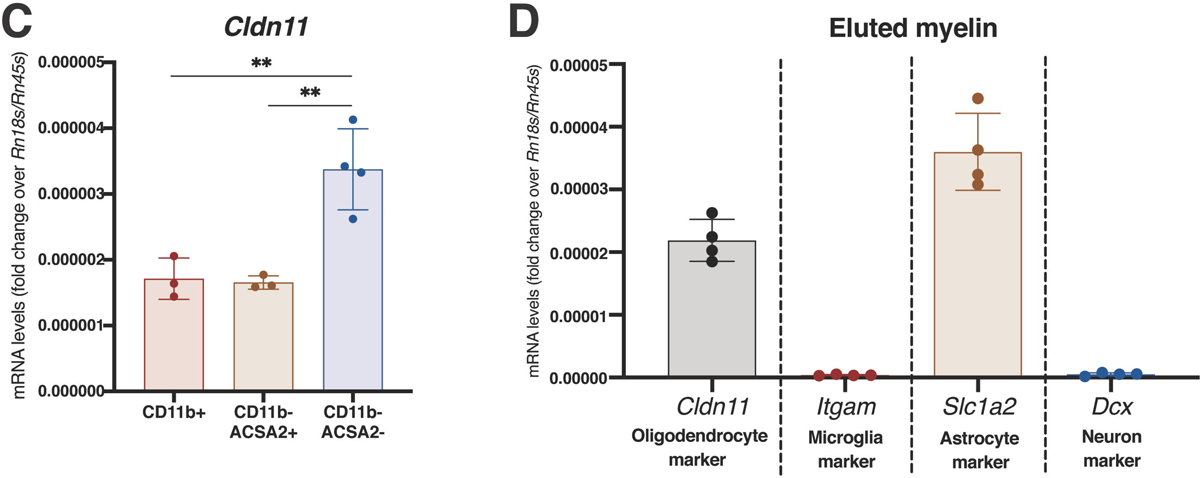
Expression levels of cell type-specific mRNA markers in sorted cell populations from whole adult mouse brain. **(A, B)** Gene expression analysis of indicated genes in sorted ACSA-2+ (astrocytes) and CD11b+ (microglia) populations (collected after removal of myelin via magnetic beads) as well as remaining cells, considered to be enriched for neurons. The threshold-crossing cycle (Ct) value for each gene was normalized by the Ct value of *Rn18s/Rn45s.* The cell populations were each collected from 4 separate mice, with each data point representing the average of 3 technical replicates (qPCR wells) from each mouse. Error bars represent standard deviation among biological replicates. Statistical differences were determined by Ordinary one-way ANOVA followed by Bonferroni’s multiple comparisons test at an ⍺=0.05 significance level; *, p<0.5; **, p<0.01; ***, p<0.001; ****, p<0.0001. (A) Organized by population. (B) Organized by marker. **(C, D)** *Cldn11* (protein: claudin-11, oligodendrocyte-specific protein, OSP) is used as an oligodendrocyte marker. (C) Gene expression analysis of *Cldn11* in Cd11b^+^ (microglia), Cd11b^-^/ACSA-2^+^ (astrocyte) and Cd11b^-^/ACSA-2^-^ (neuron) populations shows that final flow-through retains substantial *Cldn11* expression compared to other populations. (D) Gene expression analysis of indicated cell type markers in eluted myelin shows that, while *Cldn11* expression is higher than *Itgam* and *Dcx,* the myelin retains substantial expression of *S100b* (a gene known to also be enriched in oligodendrocytes) and *Slc1a2* (an astrocyte marker).

**Supplemental Figure 4:**
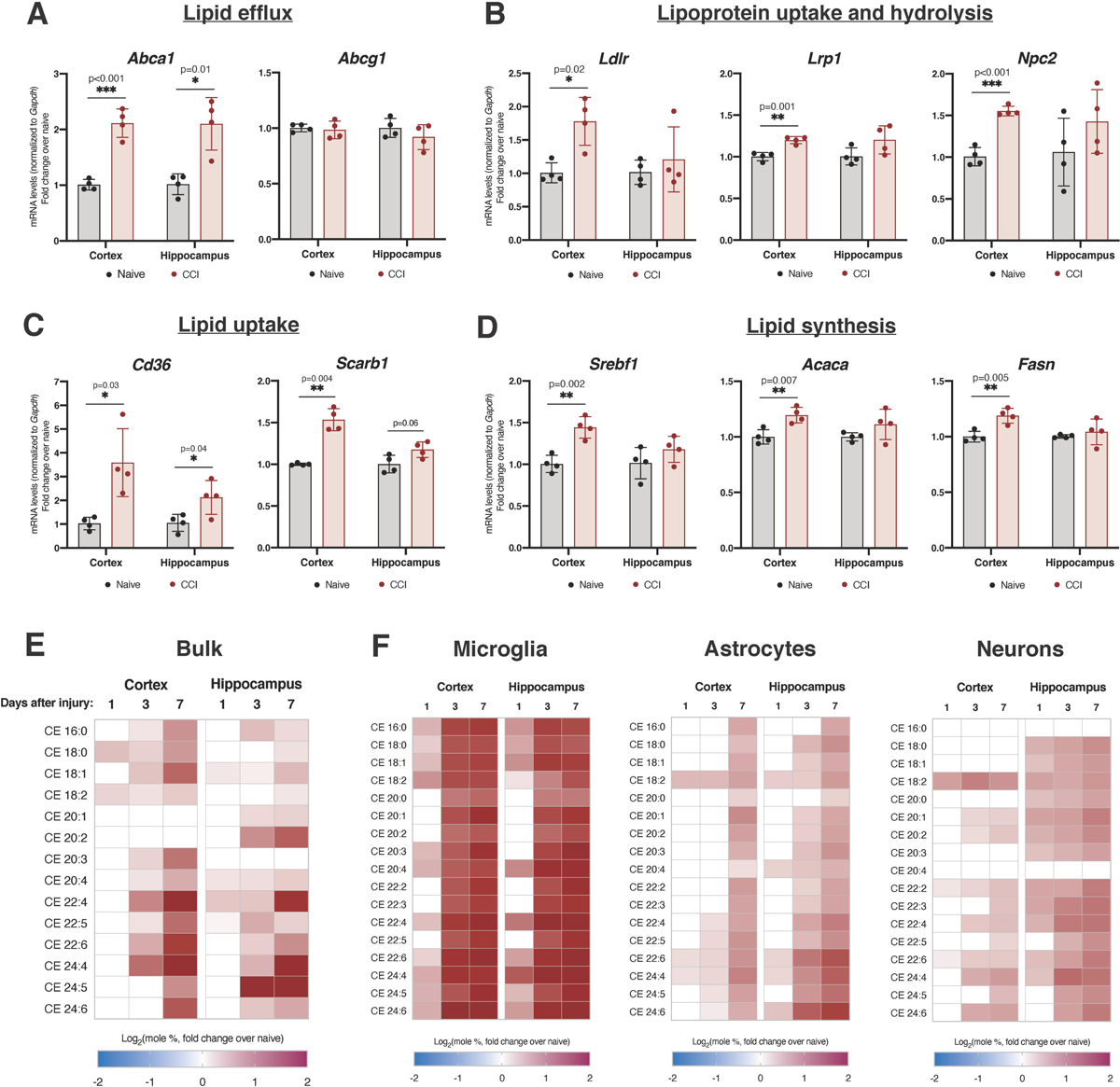
Increased lipid trafficking and cholesterol esterification in the cortex and hippocampus after CCI. **(A-D)** Expression of genes important in cellular lipid metabolism via qPCR 3d after CCI, normalized to *Gapdh*. (A) Lipid efflux transporters *Abca1* and *Abcg1*. (B) Mediators of lipoprotein uptake, *Ldlr* and *Lrp1,* and hydrolysis, *Npc2*. (C) Scavenger lipid receptors *Cd36* and *Scarb1.* (D) Regulators of *de novo* lipogenesis *Srebf1*, *Acaca* and *Fasn.* **(E)** Lipidomics levels of individual CE species in bulk ipsilateral cortical and hippocampal homogenates at multiple timepoints after CCI. This data is averaged from 3 biological replicates. **(F)** Lipidomics levels of individual CE species in sorted populations of cortical and hippocampal microglia, astrocytes and neurons at multiple timepoints after CCI. For **A-F**, statistical comparisons are made to naïve (uninjured) tissues from the same region and/or cell type harvested and assayed alongside CCI tissues (two-tailed T-test, ⍺=0.05, *, p<0.05; **, p<0.01). For **A-D,** each data point represents a separate mouse (biological replicate) and the average of 3 technical replicates (qPCR wells). Error bars represent standard deviation among biological replicates. For **E** and **F**, only statistically significant differences are represented (p<0.05).

**Supplemental Figure 5:**
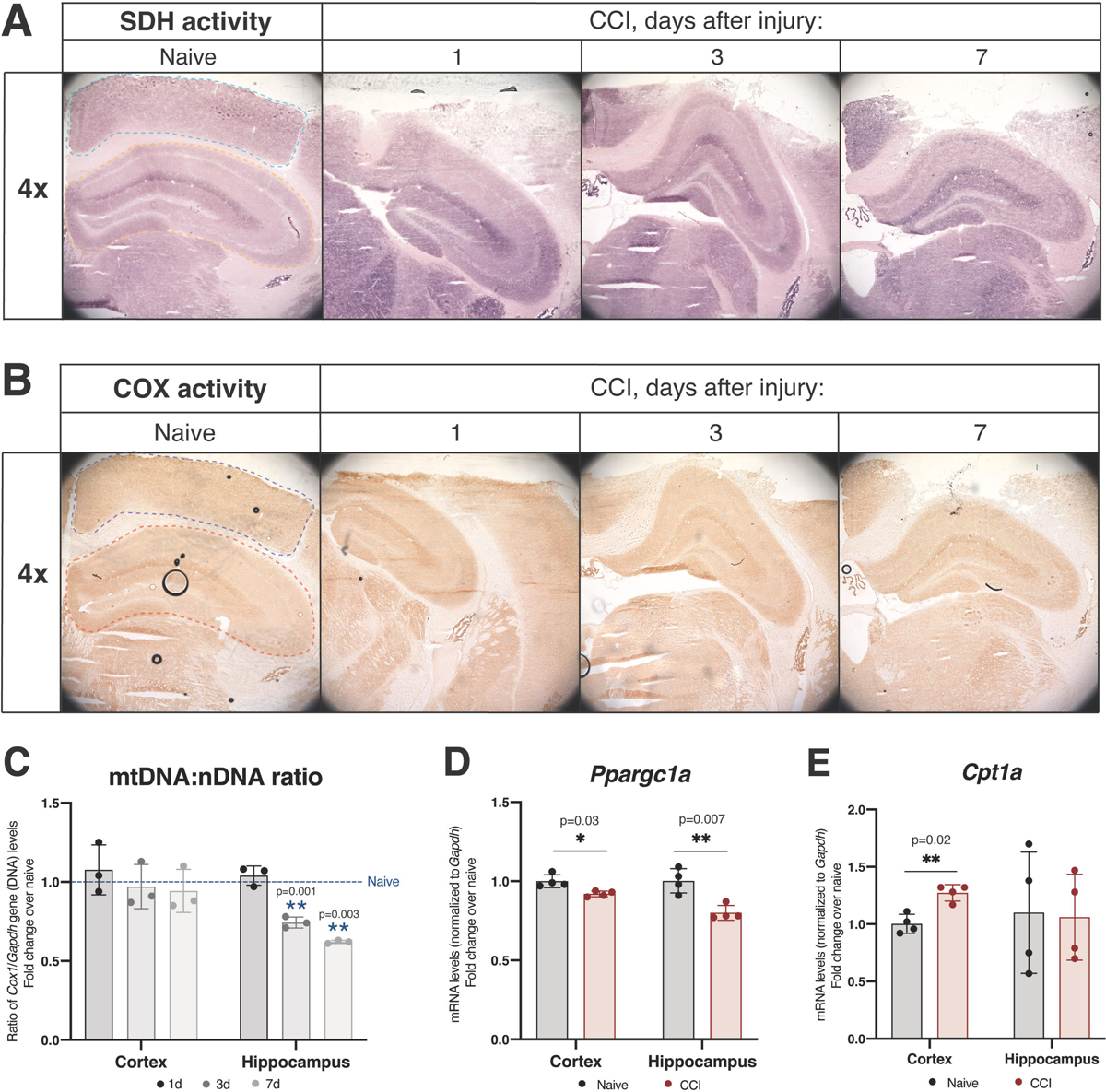
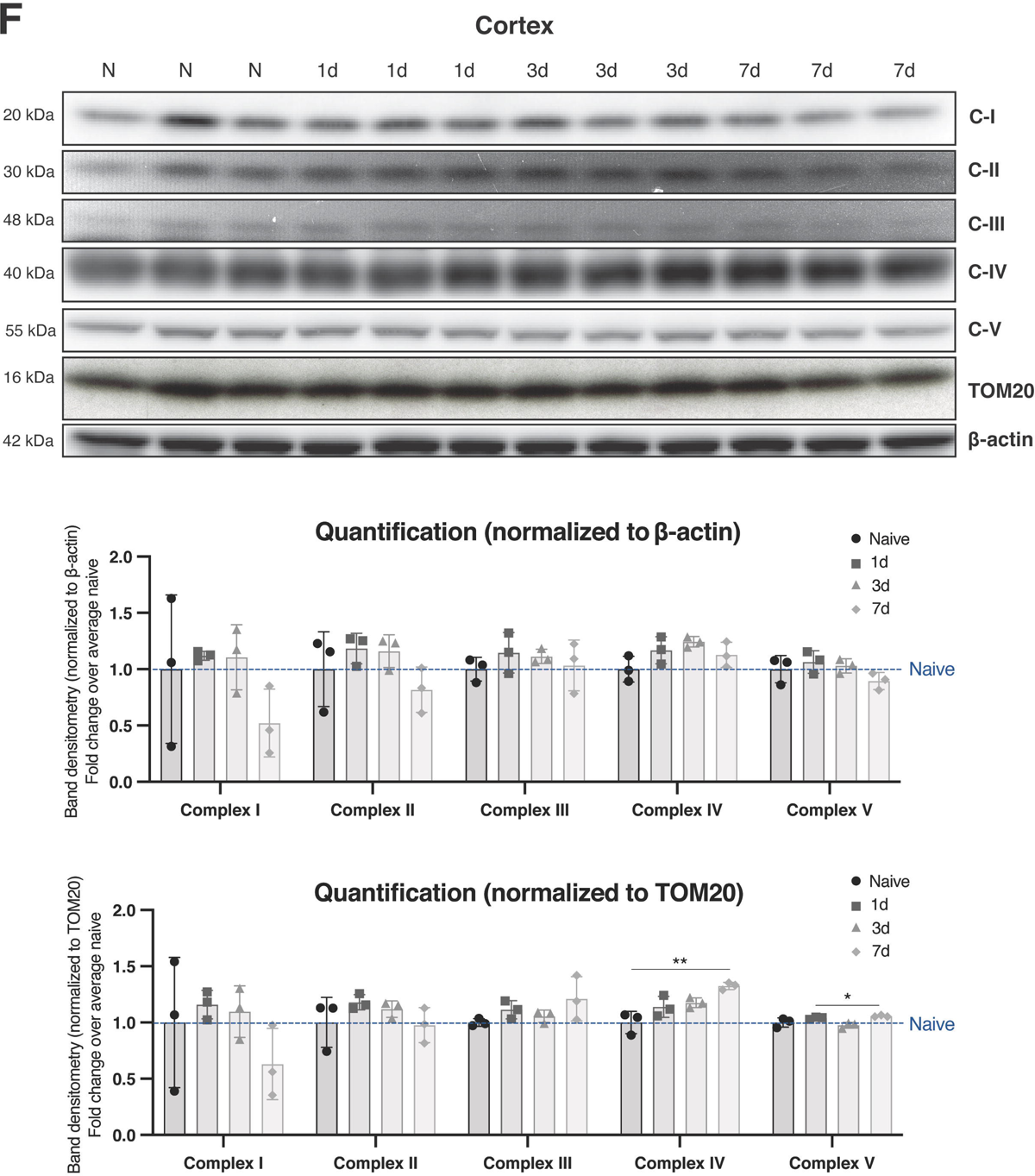
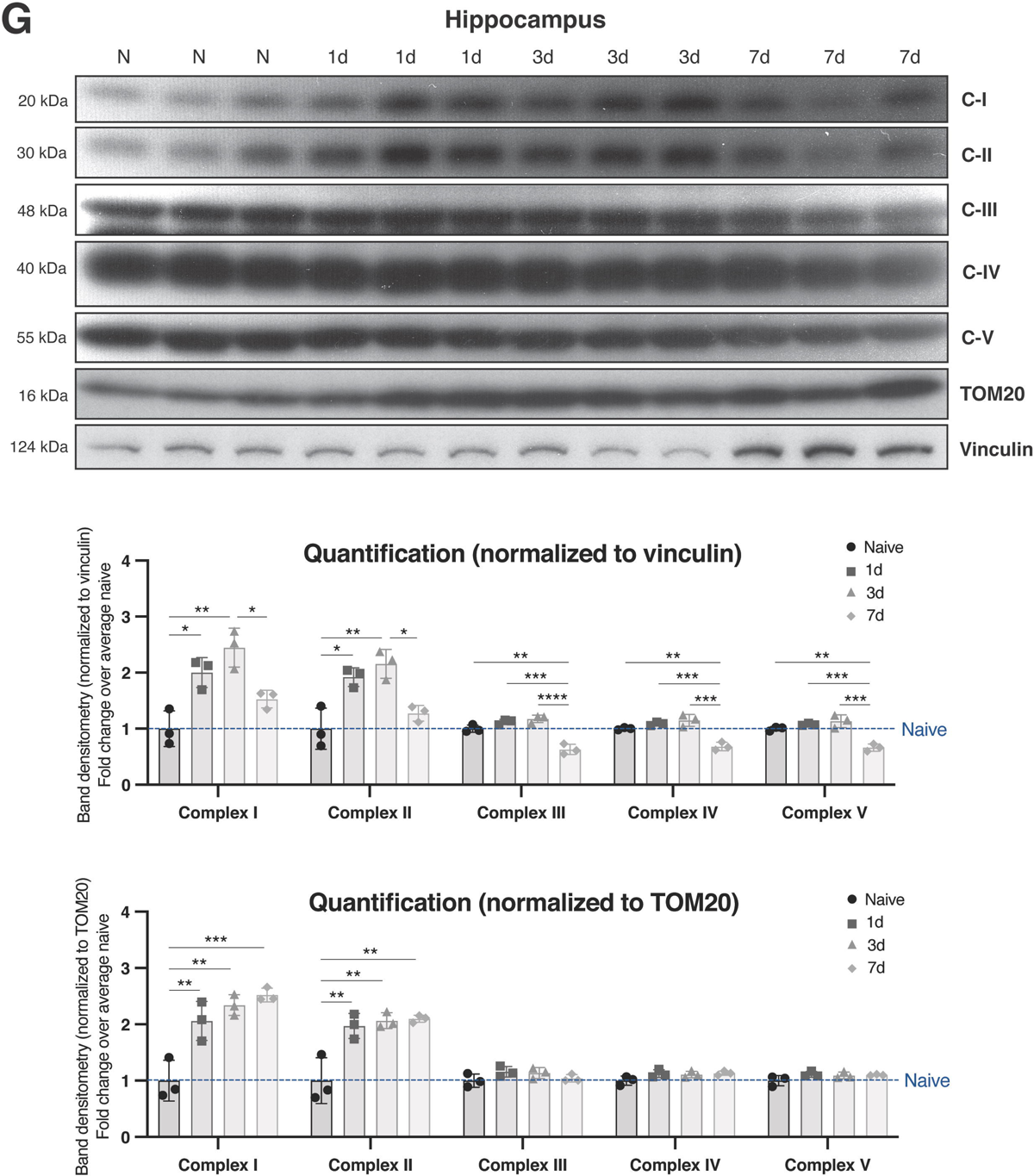
Mitochondrial regulation after CCI. **(A)** Succinate dehydrogenase (SDH, complex-II) activity staining images in the ipsilateral cortex (outlined in blue) and ipsilateral hippocampus (outlined in yellow) at multiple timepoints after CCI, representative of 3 separate mice (biological replicates) per group. **(B)** Cytochrome C oxidase (COX) activity staining images in the ipsilateral cortex (outlined in blue) and ipsilateral hippocampus (outlined in yellow) at multiple time-points after CCI, representative of 3 separate mice (biological replicates) per group. **(C)** Ratio of mitochondrial DNA (determined by *Cox1* DNA quantity) to nuclear DNA (determined by *Gapdh* DNA quantity) in the ipsilateral cortex and hippocampus at multiple timepoints after CCI. **(D)** Expression of *Ppargc1a* (encoding PGC1⍺), a regulator of mitochondrial biogenesis, via qPCR 3d after CCI. **(E)** Expression of *Cpt1a*, a regulator of mitochondrial fatty acid β-oxidation, via qPCR 3d after CCI. **(F, G)** Western blot of OxPhos complexes in both the cortex and hippocampus in naïve and CCI tissues (1, 3 and 7 days after CCI). Each lane is a separate mouse (biological replicate). Quantification is also included, with normalization to either β-actin or vinculin (whole cell markers) or TOM20 (mitochondria marker). For **C-E**, statistical comparisons are made to naïve (uninjured) tissues from the same region harvested and assayed alongside CCI tissues (two-tailed T-test; ⍺=0.05; *, p<0.05; **, p<0.01). Each data point represents a separate mouse (biological replicate) and the average of 3 technical replicates (qPCR wells). Error bars represent standard deviation among biological replicates. For **F/G**, statistical differences were determined by Ordinary one-way ANOVA followed by Bonferroni’s multiple comparisons test at an ⍺=0.05 significance level; *, p<0.5; **, p<0.01; ***, p<0.001. Error bars represent standard deviation among biological replicates.

**Supplemental Figure 6:**
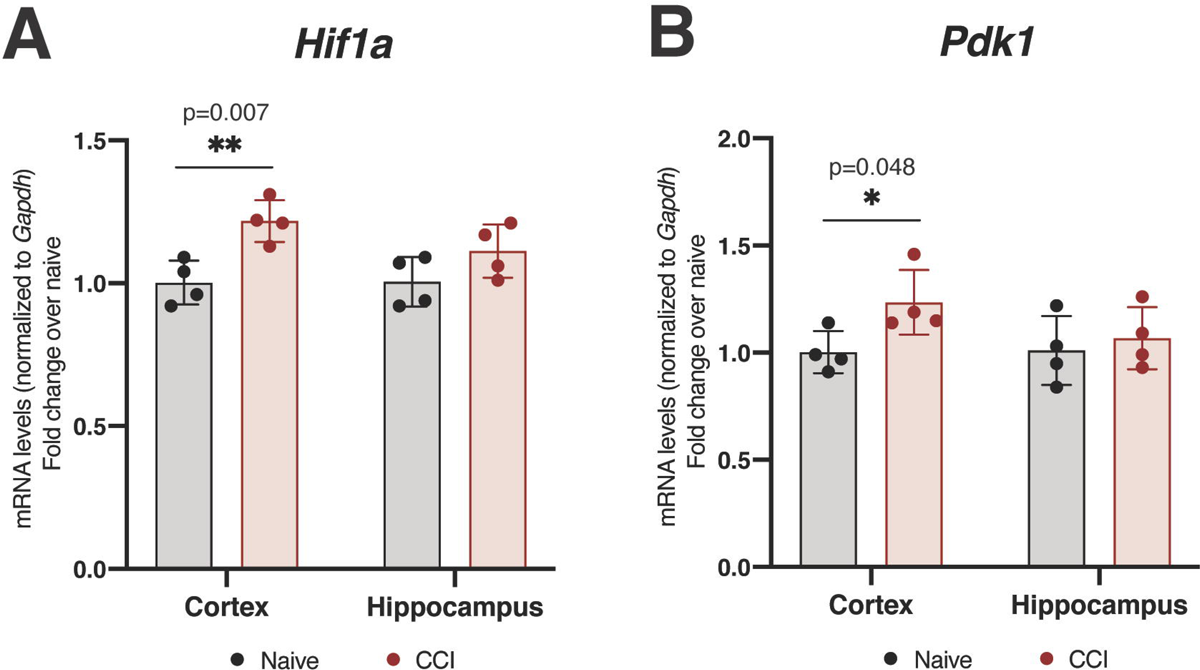
Expression of genes involved in hypoxic signaling and pyruvate regulation in the cortex and hippocampus after CCI. These qPCR analyses were conducted in ipsilateral cortical and hippocampal homogenates collected 3d after CCI. **(A)** *Hif1a*, a chief mediator of the hypoxic signaling pathway. **(B)** *Pdk1*, which encodes pyruvate dehydrogenase kinase 1. Statistical comparisons are made to naïve (uninjured) tissues from the same region harvested and assayed alongside CCI tissues (two-tailed T-test; ⍺=0.05; *, p<0.05; **, p<0.01). Each data point represents a separate mouse (biological replicate) and the average of 3 technical replicates (qPCR wells). Error bars represent standard deviation among biological replicates.

