## Supplementary material for "Alzheimer’s-associated upregulation of mitochondria-associated ER membranes after traumatic brain injury": Lipidomics source data - astrocytes

| All values represent fold change over naïve samples | | | | | | |
|---|---|---|---|---|---|---|
| **<u>Total of each lipid class:</u>** | **Ipsilateral cortex** | | | **Ipsilateral hippocampus** | | |
| **Days after injury:** | **1** | **3** | **7** | **1** | **3** | **7** |
| Free cholesterol (FC) | -0.086196 | -0.0187431 | -0.3104425 | -0.3163394 | -0.4145157 | -0.3745832 |
| Cholesteryl ester (CE) | 1.07277307 | 1.10431259 | 2.72204299 | 1.06689522 | 2.54630209 | 3.62000271 |
| CE:FC | 1.15896906 | 1.12305569 | 3.0324855 | 1.38323458 | 2.96081775 | 3.99458594 |
| Sphingomyelin (SM) | | | 1.59318762 | 1.03224506 | 0.94724311 | 0.62648455 |
| Ceramide (Cer) | | 1.00107118 | 0.77991839 | -0.3884012 | -0.3318954 | -0.433126 |
| Monohydroxylated Cer (MHCer) + Ganglioside GM3 | 0.53372077 | 0.67157117 | 1.14207201 | 1.45578661 | 1.69457523 | 0.62628733 |
| Monoglyceride (MG) | 0.29894795 | 0.66045436 | 0.87722592 | 0.64254519 | 0.56965577 | 0.63453494 |
| Diglyceride (DG) | 0.30266526 | -0.1886347 | -0.4341915 | 1.07050377 | 1.33730222 | 0.98129559 |
| Triglyceride (TG) | 0.51436791 | 0.06274974 | 0.21120728 | 1.2374797 | 0.9839552 | 1.0151136 |
| Phosphatidylcholine (PC) | | | 1.58967692 | | | 1.25200794 |
| **<u>Cholesteryl esters (CEs):</u>** | **Ipsilateral cortex** | | | **Ipsilateral hippocampus** | | |
| **Days after injury:** | **1** | **3** | **7** | **1** | **3** | **7** |
| CE 16:0 | | | 0.84620885 | | | 0.851674 |
| CE 18:0 | | | 0.64701167 | | 0.70815826 | 1.0472267 |
| CE 18:1 | | | 0.80981806 | | 0.50413758 | 0.82336967 |
| CE 18:2 | 0.67145006 | 0.66494216 | 0.89816816 | 0.49565169 | 0.51443597 | 0.95260351 |
| CE 20:0 | | | 0.44921431 | | | 0.4477186 |
| CE 20:1 | | | 1.08892979 | | 0.48377381 | 0.84043324 |
| CE 20:2 | | | 0.90822319 | | 0.48627972 | 0.89717581 |
| CE 20:3 | | | 0.7715964 | | 0.6033482 | 1.01225882 |
| CE 20:4 | | | 0.32695177 | 0.44659861 | 0.56742143 | 0.75098703 |
| CE 22:2 | | | 0.94026748 | | 0.44530145 | 0.87485509 |
| CE 22:3 | | | 0.89254686 | | 0.53541032 | 0.8761908 |
| CE 22:4 | | 0.33117232 | 0.8825332 | | 0.85765227 | 1.20550343 |
| CE 22:5 | | 0.39112399 | 1.01029503 | | 0.66840537 | 1.05873854 |
| CE 22:6 | 0.3537833 | 0.43499952 | 1.02937421 | 0.42551674 | 1.26193761 | 1.56503583 |
| CE 24:4 | 0.31654126 | 0.40929493 | 1.16137344 | 0.40133806 | 1.07189813 | 1.46073536 |
| CE 24:5 | | 0.30497456 | 0.9121443 | | 0.99027681 | 1.33557249 |
| CE 24:6 | | 0.20429757 | 0.78020582 | 0.59217516 | 1.47200294 | 1.72416325 |
| **<u>Acylcarnitines (ACs):</u>** | **Ipsilateral cortex** | | | **Ipsilateral hippocampus** | | |
| **Days after injury:** | **1** | **3** | **7** | **1** | **3** | **7** |
| AC C2:0 | | | | 1.42518629 | 2.18409884 | |
| AC C3:0 | 1.47332768 | 1.19203039 | 0.49754329 | | 1.73994166 | |
| AC C6:0 | 0.77559369 | | | | | |
| AC C12:0 | | | | | | |
| AC C14:0 | 1.13393385 | 0.84658417 | | | | |
| AC C16:0 | 1.33462669 | 0.95524928 | | 1.85707109 | 1.78445322 | 1.84997909 |
| AC C18:0 | 1.14200485 | | | 1.68859285 | 1.66975198 | 2.01634452 |
| AC C18:1 | 1.02707637 | | | | | |

| Diacylglycerols (DGs): | Ipsilateral cortex | | | Ipsilateral hippocampus | | |
|---|---|---|---|---|---|---|
| **Days after injury:** | **1** | **3** | **7** | **1** | **3** | **7** |
| DG 34:1/16:0 | 0.2003429 | 0.60267463 | 0.98853604 | 1.46554642 | 1.43227213 | 2.05482766 |
| DG 34:2/16:0 | 1.39433896 | 0.13699383 | 0.13007369 | 2.2705662 | 2.11066918 | 1.4774145 |
| DG 36:1/18:0 | 0.71918352 | 0.02986352 | -0.434983 | 1.62330718 | 1.77532489 | 1.39263499 |
| DG 36:2/18:0 | 1.27460888 | 0.24767883 | 0.00531158 | 2.29082029 | 2.00341584 | 1.32124432 |
| DG 36:2/18:1 | 0.35180771 | -0.2205572 | -0.4670022 | 1.20125041 | 1.39297382 | 1.15256811 |
| DG 36:3/18:1 | 1.29858987 | 0.21784275 | 0.06130808 | 2.21311392 | 1.51425705 | 1.52342878 |
| DG 38:2/18:1 | 0.6073977 | -0.1606457 | 0.2105065 | 1.19614303 | 1.30834366 | 0.91716878 |
| **Triacylglyerols (TGs):** | **Ipsilateral cortex** | | | **Ipsilateral hippocampus** | | |
| **Days after injury:** | **1** | **3** | **7** | **1** | **3** | **7** |
| TG 52:3/18:1 | 1.92935916 | 1.41752651 | 1.92249547 | 2.26222638 | 1.45105978 | 1.28212004 |
| TG 52:4/18:1 | 0.58294698 | 0.2116752 | 0.09706935 | 1.63628289 | 1.34314833 | 0.96280987 |
| TG 52:5/18:1 | 0.9119555 | 0.35950121 | 0.16196219 | 1.97762476 | 1.49246673 | 0.98147057 |
| TG 56:5/20:4 | 1.19805945 | 0.41324653 | 0.14595121 | 2.16082732 | 1.62497116 | 0.87477231 |
| TG 56:6/20:4 | 0.69675112 | 0.30190058 | -0.1410424 | 2.26635666 | 1.95380998 | 0.98648815 |
| TG 56:7/20:4 | 0.93913114 | 0.55418014 | -0.0627979 | 2.14142841 | 1.84849182 | 1.1163154 |
| TG 56:8/20:4 | 0.64384239 | 0.17372582 | 0.39367823 | 1.7850795 | 1.68579979 | 1.2316925 |
| TG 56:9/20:4 | 0.99737887 | 0.44781139 | 0.50964756 | 1.81106562 | 1.45098622 | 1.04094104 |
| TG 58:5/20:4 | 0.82046328 | 0.34261876 | 0.11490649 | 1.77243206 | 1.50747522 | 0.87133585 |
| TG 58:6/20:4 | 0.95304063 | 0.68557655 | 0.37626857 | 1.60989263 | 1.67582496 | 0.9991329 |
| TG 58:7/20:4 | 0.93371288 | 0.60114733 | 0.1788039 | 1.99486588 | 1.98141327 | 1.29487476 |
| TG 58:8/22:6 | 0.96375187 | 0.8324978 | 0.74935482 | 2.27001663 | 2.78415718 | 1.9109083 |
| TG 58:9/22:6 | 1.17686336 | 0.79364347 | 1.0006918 | 1.79173929 | 1.7905914 | 1.18016887 |
| TG 60:7/22:6 | 0.75594747 | 0.42535093 | 0.25344134 | 1.66407808 | 1.77296365 | 1.16133639 |
| TG 60:8/22:6 | 1.3405557 | 0.62406538 | 0.19076067 | 2.22421616 | 1.79674771 | 1.10069145 |
| TG 60:9/22:6 | 1.443239 | 0.73191805 | 0.56265362 | 1.9442133 | 1.47831189 | 0.9921153 |
