## Supplementary material for "Alzheimer’s-associated upregulation of mitochondria-associated ER membranes after traumatic brain injury": Lipidomics source data - bulk

| All values represent fold change over naïve samples | | | | | | |
|---|---|---|---|---|---|---|
| **Total of each lipid class:** | **Ipsilateral cortex** | | | **Ipsilateral hippocampus** | | |
| **Days after injury:** | **1** | **3** | **7** | **1** | **3** | **7** |
| Free cholesterol (FC) | -0.3801534 | -0.2111702 | 0.1009199 | -0.1165221 | -0.0540407 | -0.5727573 |
| Cholesteryl ester (CE) | 0.97379671 | 1.06907205 | 2.84753999 | 0.87018658 | 1.37702307 | -0.1674929 |
| CE:FC | 2.55611 | 1.75445 | 9.73742 | 1.98166 | 2.87817 | 0.16793 |
| Sphingomyelin (SM) | | 0.20029526 | 0.00845426 | | 0.25987045 | 0.63500771 |
| Ceramide (Cer) | | -0.9110107 | -1.9279894 | | -1.4312742 | -2 |
| Monohydroxylated Cer (MHCer) + Ganglioside GM3 | 0.07040329 | 0.18659694 | 0.13653355 | 0.0659479 | 0.12486797 | 0.33047825 |
| Monoglyceride (MG) | -0.1409022 | -1.2132479 | -2 | -0.2501331 | -0.1309348 | -1.4571962 |
| Diglyceride (DG) | 0.2907 | 0.15259 | -0.573 | -0.22924 | -0.3471 | 0.18873 |
| Triglyceride (TG) | 0.26186 | 0.35389 | -0.1648 | -0.68754 | -0.2824 | -0.08947 |
| Phosphatidylcholine (PC) | | 0.28494439 | 0.04707204 | | 0.09285902 | 0.16308906 |
| **Cholesteryl esters (CEs):** | **Ipsilateral cortex** | | | **Ipsilateral hippocampus** | | |
| **Days after injury:** | **1** | **3** | **7** | **1** | **3** | **7** |
| CE 16:0 | | 0.64058831 | 2.5675983 | | 1.60510653 | 0.92848555 |
| CE 18:0 | 1.5772331 | 1.15378514 | 2.41442042 | | | 0.76518745 |
| CE 18:1 | | 1.45138316 | 3.51649413 | 0.58862831 | 0.40386778 | 1.66135447 |
| CE 18:2 | 1.05241083 | 0.72097209 | 1.27838392 | | | 0.6118745 |
| CE 20:1 | | | | | 0.96027464 | 1.12348535 |
| CE 20:2 | | | | | 2.61849526 | 3.6531626 |
| CE 20:3 | | 1.09693978 | 3.19973016 | | | |
| CE 20:4 | | 0.67776103 | 1.95946106 | 0.69250999 | 0.84187207 | 1.24171604 |
| CE 22:4 | | 2.83135909 | 4.99699132 | 1.36475502 | 1.4098488 | 4.63080882 |
| CE 22:5 | | 0.71403871 | 3.39282245 | 0.25013024 | 1.97397223 | 1.14579851 |
| CE 22:6 | | 1.9886478 | 4.3753043 | | 1.20184736 | 2.64891854 |
| CE 24:4 | | 3.35064695 | 5.77316072 | | 1.81285055 | 4.93269545 |
| CE 24:5 | | | 2.67111576 | | 8.66654279 | 9.15296587 |
| CE 24:6 | | | 3.85559898 | | 1.38806623 | 2.07021375 |
| **Acylcarnitines (ACs):** | **Ipsilateral cortex** | | | **Ipsilateral hippocampus** | | |
| **Days after injury:** | **1** | **3** | **7** | **1** | **3** | **7** |
| AC C2:0 | 1.6580391 | 4.8360782 | 3.89844279 | | 0.62769656 | 2.64441332 |
| AC C3:0 | 0.8675618 | 2.89920911 | 2.4422593 | | 0.91259936 | 3.01777261 |
| AC C6:0 | | 1.24336182 | 1.35818472 | 0.77152344 | 0.65179884 | 1.07935552 |
| AC C12:0 | | | 1.93646988 | | | |
| AC C14:0 | | | 1.83652217 | | | |
| AC C16:0 | | | 3.08405693 | | 0.74873488 | 1.49351121 |
| AC C18:0 | | | | | | |
| AC C18:1 | | 1.01139736 | 2.54930851 | | 0.78190039 | 1.36295102 |

| **Diacylglycerols (DGs):** | **Ipsilateral cortex** | | | **Ipsilateral hippocampus** | | |
|---|---|---|---|---|---|---|
| **Days after injury:** | **1** | **3** | **7** | **1** | **3** | **7** |
| DG 30:0/14:0 | | | | | | |
| DG 30:1/14:0 | | | | | | |
| DG 32:0/16:0 | | | | | | |
| DG 32:1/16:0 | | 1.09591213 | 1.10396008 | | | |
| DG 32:2/16:1 | | | | | 0.51014549 | 0.97563145 |
| DG 34:0/16:0 | | | | | | |
| DG 34:1/16:0 | | | | | | |
| DG 34:2/16:0 | | 0.917404 | 1.10302385 | | 0.97696106 | 0.08298899 |
| DG 34:2/16:1 | | | | | | |
| DG 36:0/18:0 | | | | | | |
| DG 36:1/18:0 | | 0.5286135 | 0.85356029 | 0.40842396 | -0.2302126 | -1.1645944 |
| DG 36:2/18:0 | | | | | | -0.5121336 |
| DG 36:2/18:1 | | | | | | -0.9280278 |
| DG 36:3/18:1 | | | | | | -0.3634279 |
| DG 36:4/18:0 | | | | | | 0.92727969 |
| DG 38:1/18:0 | | 0.57277956 | 1.30753876 | 0.6164709 | -0.6570594 | -1.1148859 |
| DG 38:2/18:0 | | 0.49355671 | 1.41771995 | 0.21686686 | 0.84049646 | 1.10577875 |
| DG 38:2/18:1 | | 0.50731603 | 1.04492957 | 0.3088645 | -0.4083384 | -0.7959325 |
| DG 38:3/18:0 | | 0.75815236 | 0.8979186 | | 0.39499789 | 1.24771764 |
| DG 38:3/18:1 | | | | | | |
| DG 38:4/18:0 | | | 0.916892 | | | |
| DG 38:4/18:1 | | 1.13102664 | 1.04881262 | | | |
| DG 40:4/18:0 | | | | | | |
| DG 40:4/18:1 | | | | | | |
| DG 40:5/18:0 | | | | | | |
| DG 40:5/18:1 | | | 1.17267433 | | | -0.9633568 |
| DG 40:6/18:0 | | | | -0.3239013 | -1.2845674 | -2.401739 |
| DG 40:6/18:1 | | | | | | |

| **Triacylglyerols (TGs):** | **Ipsilateral cortex** | | | **Ipsilateral hippocampus** | | |
|---|---|---|---|---|---|---|
| **Days after injury:** | **1** | **3** | **7** | **1** | **3** | **7** |
| TG 48:0/16:0 | | | | | | |
| TG 48:1/16:0 | | | | | | |
| TG 50:0/16:0 | | | | | | |
| TG 50:1/16:1 | | | | | | |
| TG 50:2/16:1 | | | | | | |
| TG 50:3/16:1 | | | | | 0.35775125 | 1.10055947 |
| TG 52:0/18:0 | | | | | | |
| TG 52:1/18:0 | | | | | | |
| TG 52:2/18:0 | | | | | | |
| TG 52:3/18:1 | | | | | 0.26864348 | 0.93111873 |
| TG 52:4/18:1 | | | 0.62450061 | | | |
| TG 52:5/18:1 | | | 0.99682841 | 1.34528476 | 1.31283846 | 2.27633601 |
| TG 52:5/20:4 | | | | | | |
| TG 54:0/18:0 | | | | | | |
| TG 54:1/18:0 | | | | | | |
| TG 54:2/18:0 | | | | -0.2841957 | 0.61890902 | 0.95758029 |
| TG 54:3/18:0 | | | 0.94452972 | -0.6378727 | 0.51389424 | 1.28280789 |
| TG 54:4/18:1 | | 0.47986003 | 0.43699278 | -0.4133076 | 0.08368501 | 1.38791549 |
| TG 54:4/20:4 | | 0.29078126 | 0.70980231 | | | |
| TG 54:5/18:1 | 0.36294474 | 0.58582115 | 0.65555745 | | | 0.92901956 |
| TG 54:5/20:4 | 0.12391231 | 0.20421286 | 0.42813631 | | | |
| TG 54:6/18:1 | | | | | 1.39563297 | 2.93472613 |
| TG 54:6/20:4 | | | | | | |
| TG 54:7/18:1 | | | | | | 1.67507775 |
| TG 54:7/20:4 | 0.92686946 | 1.96355582 | 2.03495495 | | | 1.18504232 |
| TG 56:3/18:1 | 0.45992744 | 0.79521832 | 1.28101612 | | 0.25647864 | 0.90008933 |
| TG 56:4/18:1 | | | | | | |
| TG 56:4/20:4 | | | | | | |
| TG 56:5/18:1 | | | | 0.15764351 | 0.24676484 | 0.7462881 |
| TG 56:5/20:4 | | | | | | |
| TG 56:6/20:4 | | | | 0.08841455 | 0.23852648 | -0.2245801 |
| TG 56:7/20:4 | 1.21551795 | 1.29763885 | 0.92462438 | 0.04250137 | 0.29197159 | -0.7721246 |
| TG 56:8/20:4 | | | | 0.11910416 | 0.55564913 | -0.8713653 |
| TG 56:9/20:4 | | | | 1.60818792 | 0.98228877 | -0.4615431 |
| TG 58:5/20:4 | | | | 1.35770441 | 1.16248782 | 1.35866422 |
| TG 58:6/20:4 | | | | | | |
| TG 58:7/20:4 | | | 0.83456128 | | | |
| TG 58:8/22:6 | 0.60205184 | 0.92326177 | 0.67946142 | | | |
| TG 58:9/22:6 | | | | | | |
| TG 60:7/22:6 | | | | | | 2.24821425 |
| TG 60:8/22:6 | 0.81793391 | 1.07254244 | | | | 2.19894177 |
| TG 60:9/22:6 | 1.11873031 | 1.86802279 | 1.7371794 | | | 1.82897616 |
