## Supplementary material for "Alzheimer’s-associated upregulation of mitochondria-associated ER membranes after traumatic brain injury": Lipidomics source data - microglia

| All values represent fold change over naïve samples | | | | | | |
|---|---|---|---|---|---|---|
| **Total of each lipid class:** | **Ipsilateral cortex** | | | **Ipsilateral hippocampus** | | |
| **Days after injury:** | **1** | **3** | **7** | **1** | **3** | **7** |
| Free cholesterol (FC) | 0.14882428 | -0.3393934 | -0.5596058 | -0.5454888 | -0.7832872 | -1.1996666 |
| Cholesteryl ester (CE) | 2.37821436 | 5.52611352 | 6.13258771 | 3.29181441 | 5.94338866 | 6.37561505 |
| CE:FC | 2.22939008 | 5.86550689 | 6.6921935 | 3.8373032 | 6.72667583 | 7.57528165 |
| Sphingomyelin (SM) | | -0.2953701 | -0.7239071 | -0.1166636 | -0.6858312 | |
| Ceramide (Cer) | | 0.34859592 | 0.80394934 | 0.56231051 | 0.82444946 | |
| Monohydroxylated Cer (MHCer) + Ganglioside GM3 | 0.51674634 | 0.26063861 | -0.4208551 | 1.37141356 | 1.44645861 | 1.23732236 |
| Monoglyceride (MG) | -0.0712384 | -0.2917312 | -0.86805 | | -0.5719425 | -0.9919413 |
| Diglyceride (DG) | 0.79194689 | 0.83115712 | 0.05629368 | 0.49049908 | -0.4313938 | -0.8159418 |
| Triglyceride (TG) | 2.22272869 | 1.67885346 | -0.1628497 | 1.53679084 | 0.51271121 | -0.2900744 |
| Phosphatidylcholine (PC) | | | | | | |
| **Cholesteryl esters (CEs):** | **Ipsilateral cortex** | | | **Ipsilateral hippocampus** | | |
| **Days after injury:** | **1** | **3** | **7** | **1** | **3** | **7** |
| CE 16:0 | 0.72741183 | 1.69983929 | 1.76703713 | 0.9986088 | 1.71661132 | 1.65104764 |
| CE 18:0 | 0.48906551 | 1.56554078 | 1.6112384 | 0.8389311 | 1.59333822 | 1.49678947 |
| CE 18:1 | 0.64731521 | 1.63261838 | 1.70310478 | 1.04141946 | 1.76507519 | 1.73679998 |
| CE 18:2 | 1.02600775 | 1.62877892 | 1.64749514 | 0.27261587 | 1.14146296 | 1.5791201 |
| CE 20:0 | | 1.25501535 | 1.39471723 | | 1.21524296 | 1.23131245 |
| CE 20:1 | | 1.6024512 | 1.92672695 | | 1.64818047 | 1.80505781 |
| CE 20:2 | | 1.36012821 | 1.52579082 | | 1.38557374 | 1.63290822 |
| CE 20:3 | 0.69103041 | 1.50647431 | 1.88447602 | | 1.67846044 | 2.01948569 |
| CE 20:4 | 0.74144348 | 1.48008893 | 1.65335428 | 1.10371536 | 1.89704732 | 1.93983179 |
| CE 22:2 | | 1.54257244 | 1.80822252 | | 1.59763161 | 1.91735424 |
| CE 22:3 | | 1.51826282 | 1.73456788 | | 1.69297858 | 1.8213938 |
| CE 22:4 | 0.62215888 | 1.78934559 | 1.96542462 | 1.26182883 | 2.29429717 | 2.33821274 |
| CE 22:5 | | 1.53877183 | 1.79673932 | | 1.54121878 | 1.86300251 |
| CE 22:6 | 0.45492299 | 1.86504761 | 2.07878627 | 0.91339066 | 1.86191621 | 2.12920916 |
| CE 24:4 | 0.79577063 | 1.8666931 | 1.84767205 | 1.50076361 | 2.44127283 | 2.39631416 |
| CE 24:5 | 0.74519034 | 1.56202379 | 1.89660679 | 0.69114342 | 1.59197287 | 1.89503889 |
| CE 24:6 | 0.78112457 | 2.03072037 | 2.1661341 | 0.94922221 | 1.91688582 | 2.10502989 |
| **Acylcarnitines (ACs):** | **Ipsilateral cortex** | | | **Ipsilateral hippocampus** | | |
| **Days after injury:** | **1** | **3** | **7** | **1** | **3** | **7** |
| AC C2:0 | 1.55000774 | 2.99846189 | 1.06457635 | 0.68046245 | 1.81512291 | 1.52032452 |
| AC C3:0 | | | | | | 0.62874415 |
| AC C6:0 | 0 | 0 | 0 | 0 | 0 | 0 |
| AC C12:0 | | | | | | |
| AC C14:0 | 1.12691563 | 2.50656947 | 1.52473291 | | | |
| AC C16:0 | 1.59564502 | 2.60502356 | 1.97038387 | 1.10360512 | 2.30796579 | 1.44464296 |
| AC C18:0 | 2.01462447 | 3.77645348 | 2.40805167 | 1.50308326 | 2.80450239 | 1.6918076 |
| AC C18:1 | 1.35879074 | 2.70898299 | 1.5457758 | | | |

| Diacylglycerols (DGs): | Ipsilateral cortex | | | Ipsilateral hippocampus | | |
|---|---|---|---|---|---|---|
| **Days after injury:** | **1** | **3** | **7** | **1** | **3** | **7** |
| DG 34:1/16:0 | | | 0.30572944 | 0.34244331 | -0.802407 | -1.1548327 |
| DG 34:2/16:0 | 2.34091917 | 2.32348674 | 0.49567932 | 1.73803845 | 0.16869319 | -0.4218023 |
| DG 36:1/18:0 | 1.24274753 | 1.51213543 | 0.16808331 | 1.32631409 | 0.60581712 | -0.4469639 |
| DG 36:2/18:0 | 2.18842851 | 2.29692066 | 0.36327966 | 1.94181701 | 0.39579723 | -0.4122433 |
| DG 36:2/18:1 | 1.14257081 | 1.07521234 | -0.2377663 | 0.39498977 | -0.7195493 | -1.6138015 |
| DG 36:3/18:1 | 3.084278 | 3.2417028 | 0.90724216 | 1.17177465 | -0.2487866 | -0.9441931 |
| DG 38:2/18:1 | 0.28384783 | 0.08667362 | -1.0145405 | 0.1595029 | -1.4817857 | -1.2608 |
| **Triacylglyerols (TGs):** | **Ipsilateral cortex** | | | **Ipsilateral hippocampus** | | |
| **Days after injury:** | **1** | **3** | **7** | **1** | **3** | **7** |
| TG 52:3/18:1 | 1.38944952 | 1.17673918 | | 1.12984278 | | 0.29595884 |
| TG 52:4/18:1 | 3.06202396 | 3.11551846 | | 2.37800117 | 1.46396164 | -0.684067 |
| TG 52:5/18:1 | 3.57314967 | 3.01767105 | | 3.14381103 | 1.31933279 | -0.535074 |
| TG 56:5/20:4 | 3.3203921 | 2.81826915 | | 3.17544658 | 1.00140518 | -0.3378561 |
| TG 56:6/20:4 | 2.34949677 | 1.67083023 | | 2.59966969 | 0.85327734 | -0.5021159 |
| TG 56:7/20:4 | 2.96986106 | 2.13497208 | | 3.17522512 | 0.97668188 | -0.536131 |
| TG 56:8/20:4 | 2.39946947 | 3.34633936 | | 1.6716423 | 1.21806198 | -0.4551967 |
| TG 56:9/20:4 | 2.91591829 | 3.35322001 | 0.59097671 | 2.31953521 | 1.37332087 | -0.404448 |
| TG 58:5/20:4 | 2.4657819 | 2.70157299 | -0.0123327 | 2.2753614 | 1.10296921 | -0.5293892 |
| TG 58:6/20:4 | 4.02374992 | 4.34802696 | 0.20721576 | 3.04395652 | 2.3433031 | -0.5083695 |
| TG 58:7/20:4 | 3.17385096 | 3.72300176 | 0.02729351 | 2.99897563 | 1.84415383 | -0.3703041 |
| TG 58:8/22:6 | 3.30564242 | 5.20531501 | 2.55275781 | 2.14707519 | 2.51266125 | 0.32020035 |
| TG 58:9/22:6 | 2.99593152 | 4.38375227 | 1.22546943 | 4.62155355 | 4.31632135 | 2.09977606 |
| TG 60:7/22:6 | 2.81428591 | 3.75575605 | 0.39407169 | 2.33059568 | 1.76397993 | -0.6156973 |
| TG 60:8/22:6 | 2.94713863 | 2.37898357 | -0.2358589 | 2.29772533 | 0.51710543 | -0.8421006 |
| TG 60:9/22:6 | 3.23374328 | 2.61303633 | -0.1455517 | 2.79053528 | 0.64573794 | -0.6386789 |
