## Supplementary material for "Alzheimer’s-associated upregulation of mitochondria-associated ER membranes after traumatic brain injury": Lipidomics source data - neurons

| | | | | | | |
|---|---|---|---|---|---|---|
| All values represent fold change over naïve samples | | | | | | |
| **Total of each lipid class:** | **Ipsilateral cortex** | | | **Ipsilateral hippocampus** | | |
| **Days after injury:** | **1** | **3** | **7** | **1** | **3** | **7** |
| Free cholesterol (FC) | -0.0474476 | 0.03493972 | -0.0956349 | -0.9693679 | -0.9336163 | -0.663681 |
| Cholesteryl ester (CE) | 0.44029786 | 1.15977684 | 1.17411 | 1.95023011 | 2.41059532 | 3.02959146 |
| CE:FC | 0.48774544 | 1.12483712 | 1.26974494 | 2.91959805 | 3.34421165 | 3.69327247 |
| Sphingomyelin (SM) | | 0.86757438 | 1.66149552 | 1.5301011 | 1.48242568 | 1.22010011 |
| Ceramide (Cer) | | 0.95423849 | 0.84922179 | -1.7646748 | -1.7468525 | -1.3109063 |
| Monohydroxylated Cer (MHCer) + Ganglioside GM3 | -0.0960119 | -0.4016752 | -0.5138069 | 1.58421169 | 1.62036136 | 0.73982214 |
| Monoglyceride (MG) | -0.0105085 | -0.353371 | 0.86457882 | 0.94728445 | 1.11237797 | 1.42916884 |
| Diglyceride (DG) | -0.2861985 | -0.1000616 | -0.6872579 | 0.03742771 | -0.1140908 | -0.1434709 |
| Triglyceride (TG) | -0.1199492 | -0.201325 | -0.1968648 | 0.68874447 | 0.94254442 | 0.14437927 |
| Phosphatidylcholine (PC) | | 0.81702399 | 1.6209849 | | 3.07162762 | 2.61006903 |
| **Cholesteryl esters (CEs):** | **Ipsilateral cortex** | | | **Ipsilateral hippocampus** | | |
| **Days after injury:** | **1** | **3** | **7** | **1** | **3** | **7** |
| CE 16:0 | | | | | | |
| CE 18:0 | | | | 0.75881629 | 0.8892578 | 1.05003012 |
| CE 18:1 | | | | 0.61067952 | 0.83655048 | 0.95981708 |
| CE 18:2 | 0.91978839 | 1.17721873 | 1.00157912 | 0.8945703 | 0.83287103 | 0.99082069 |
| CE 20:0 | | | | 0.6268618 | 0.72855395 | 0.89043997 |
| CE 20:1 | | 0.4159602 | 0.48024051 | 0.89194381 | 0.92143612 | 1.18334193 |
| CE 20:2 | | 0.358515 | 0.56307661 | 0.75842998 | 0.87622638 | 1.09323138 |
| CE 20:3 | | | | 0.61099416 | 0.73318058 | 0.91003941 |
| CE 20:4 | | | | | | |
| CE 22:2 | 0.24748835 | 0.42932732 | 0.61815364 | 0.85607252 | 0.85401146 | 1.21031677 |
| CE 22:3 | | | 0.59947948 | 0.85572513 | 1.15090558 | 1.28482414 |
| CE 22:4 | | 0.54755734 | 0.636468 | 0.73914093 | 1.24149972 | 1.27576327 |
| CE 22:5 | | | 0.42772617 | | 0.75777629 | 1.05255505 |
| CE 22:6 | 0.18971624 | 0.52290524 | 0.53876604 | 0.63910648 | 0.92276254 | 1.22124284 |
| CE 24:4 | | 0.78877595 | 0.94626541 | 0.58844241 | 1.36424152 | 1.27935247 |
| CE 24:5 | | | 0.66419675 | | 0.91363349 | 1.09388683 |
| CE 24:6 | | 0.81913351 | 0.89739995 | | 0.91851936 | 1.23344799 |
| **Acylcarnitines (ACs):** | **Ipsilateral cortex** | | | **Ipsilateral hippocampus** | | |
| **Days after injury:** | **1** | **3** | **7** | **1** | **3** | **7** |
| AC C2:0 | | | 1.40430846 | | 0.70700889 | 1.67642118 |
| AC C3:0 | 0.88877829 | 1.09561686 | 1.20967309 | | | 1.71340506 |
| AC C6:0 | 1.09516545 | 1.29174059 | 1.59740678 | 0.92483645 | 1.53396974 | 2.47444065 |
| AC C12:0 | | | | | | |
| AC C14:0 | | 1.31747871 | | | 0.7200373 | |
| AC C16:0 | 1.07913015 | 1.53707637 | | | 1.93300759 | |
| AC C18:0 | 0.90406098 | 1.49143797 | | | 1.91885807 | 1.25346136 |
| AC C18:1 | 0.95947162 | 1.26567008 | | | 1.25224413 | |

| Diacylglycerols (DGs): | Ipsilateral cortex | | | Ipsilateral hippocampus | | |
|---|---|---|---|---|---|---|
| Days after injury: | 1 | 3 | 7 | 1 | 3 | 7 |
| DG 34:1/16:0 | 0.00812566 | -0.1425053 | -0.1439297 | 1.09439534 | 0.88237304 | 1.32223296 |
| DG 34:2/16:0 | 0.50713317 | 0.29890753 | -0.1581742 | 1.82794085 | 1.25235601 | 1.06462897 |
| DG 36:1/18:0 | -0.0347053 | 0.27513336 | -0.7168712 | 1.04298159 | 0.62562672 | 0.41253359 |
| DG 36:2/18:0 | 0.561848 | 0.33569167 | -0.0235304 | 2.03295572 | 1.38232882 | 1.07411462 |
| DG 36:2/18:1 | -0.118639 | 0.05150681 | -0.3994335 | 0.70360068 | 0.57322713 | 0.39277698 |
| DG 36:3/18:1 | 0.61651363 | 0.11356977 | -0.0427577 | 1.87730204 | 1.22379841 | 1.00195695 |
| DG 38:2/18:1 | 0.09550208 | -0.0449832 | -0.137919 | 1.74432226 | 1.34602238 | 1.80190843 |
| Triacylglyerols (TGs): | Ipsilateral cortex | | | Ipsilateral hippocampus | | |
| Days after injury: | 1 | 3 | 7 | 1 | 3 | 7 |
| TG 52:3/18:1 | 1.73674837 | 1.35643267 | 1.29399409 | 3.05820886 | 3.60253618 | 3.19714445 |
| TG 52:4/18:1 | -0.0674054 | -0.1291138 | -0.2538524 | 1.32102483 | 1.44193799 | 0.36302184 |
| TG 52:5/18:1 | 0.19431032 | 0.19656493 | 0.25545047 | 1.91578521 | 2.0487853 | 0.97773608 |
| TG 56:5/20:4 | 0.21201438 | 0.08050535 | 0.21959166 | 1.71956933 | 1.99611798 | 0.76952527 |
| TG 56:6/20:4 | -0.1733944 | 0.08822731 | -0.0902261 | 1.57108827 | 2.04357373 | 0.72301924 |
| TG 56:7/20:4 | 0.05774702 | 0.2693862 | 0.3694244 | 1.68724411 | 1.98920106 | 0.58196543 |
| TG 56:8/20:4 | 0.34058091 | -0.0509507 | 0.00078129 | 1.9586548 | 1.96995779 | 0.77437048 |
| TG 56:9/20:4 | 0.27936082 | 0.06539186 | 0.10141681 | 2.21522987 | 2.03610416 | 1.2017785 |
| TG 58:5/20:4 | 0.13310952 | 0.05314384 | 0.20271654 | 1.71727619 | 1.94706526 | 0.91868626 |
| TG 58:6/20:4 | 0.23997068 | 0.33787932 | 0.05546828 | 1.91149267 | 2.02080016 | 0.71297127 |
| TG 58:7/20:4 | 0.16191285 | 0.25488739 | 0.04363859 | 1.57733068 | 1.74174102 | 0.63358797 |
| TG 58:8/22:6 | 0.2888732 | 0.42021954 | -0.0758481 | 1.87710849 | 2.11663576 | 0.77723568 |
| TG 58:9/22:6 | 0.37445317 | 0.33034939 | 0.12358472 | 2.16704019 | 2.05876892 | 0.95047647 |
| TG 60:7/22:6 | 0.08850003 | 0.02547042 | -0.0403956 | 1.54448337 | 1.70626719 | 0.53919557 |
| TG 60:8/22:6 | 0.44841176 | 0.44624632 | 0.36396066 | 1.77554599 | 2.12270567 | 1.01335374 |
| TG 60:9/22:6 | 0.33430853 | 0.21464164 | 0.17353032 | 1.96377516 | 2.0645275 | 1.07419632 |
